# Target-conditioned diffusion generates potent TNFR superfamily antagonists and agonists

**DOI:** 10.1101/2024.09.13.612773

**Authors:** Matthias Glögl, Aditya Krishnakumar, Robert J. Ragotte, Inna Goreshnik, Brian Coventry, Asim K. Bera, Alex Kang, Emily Joyce, Green Ahn, Buwei Huang, Wei Yang, Wei Chen, Mariana Garcia Sanchez, Brian Koepnick, David Baker

**Affiliations:** Department of Biochemistry, University of Washington, Seattle, WA, USA; Institute for Protein Design, University of Washington, Seattle, WA, USA; Howard Hughes Medical Institute, University of Washington, Seattle, WA, USA

## Abstract

Despite progress in designing protein binding proteins, the shape matching of designs to targets is lower than in many native protein complexes, and design efforts have failed for TNF receptor (TNFR1) and other protein targets with relatively flat and polar surfaces. We hypothesized that free diffusion from random noise could generate shape-matched binders for challenging targets, and tested this on TNFR1. We obtain designs with low picomolar affinity whose specificity can be completely switched to other family members using partial diffusion. Designs function as antagonists or as superagonists when presented at higher valency for OX40 and 4-1BB. The ability to design high-affinity and specificity antagonists and agonists for pharmacologically important targets in silico presages a new era in which binders are made by computation rather than immunization or random screening approaches.

## Introduction

The design of proteins that bind with high affinity and specificity to targets of interest is a long-standing challenge in computational structural biology with applications in therapeutics, diagnostics, and beyond(*1, 2*). To address this problem, protein design methods have generally relied on pre-existing sets of scaffolds—either native proteins or de novo designs—with well defined tertiary structures (*3–6*). For example, binders were generated for a number of targets by docking with a large set of idealized ∼65 residue protein scaffolds and identifying those capable of hosting the lowest energy binding interactions (*3*). RFdiffusion was used to generate binders conditioned on similar ideal scaffolds; unconstrained diffusion was also tested but only yielded simple three or four helix bundles (*7*). The advantage of using small ideal scaffolds with regular secondary structure elements and packing is that following sequence design, a reasonable fraction of designs are likely to fold as expected, but this limits the extent of shape matching achievable, particularly for targets with relatively flat surfaces lacking concavities for small miniproteins to fit into. Indeed, the contact molecular surface (CMS) of de novo-designed binding proteins to date is lower than that of many native protein complexes (Fig. 1A).

**Fig. 1.**
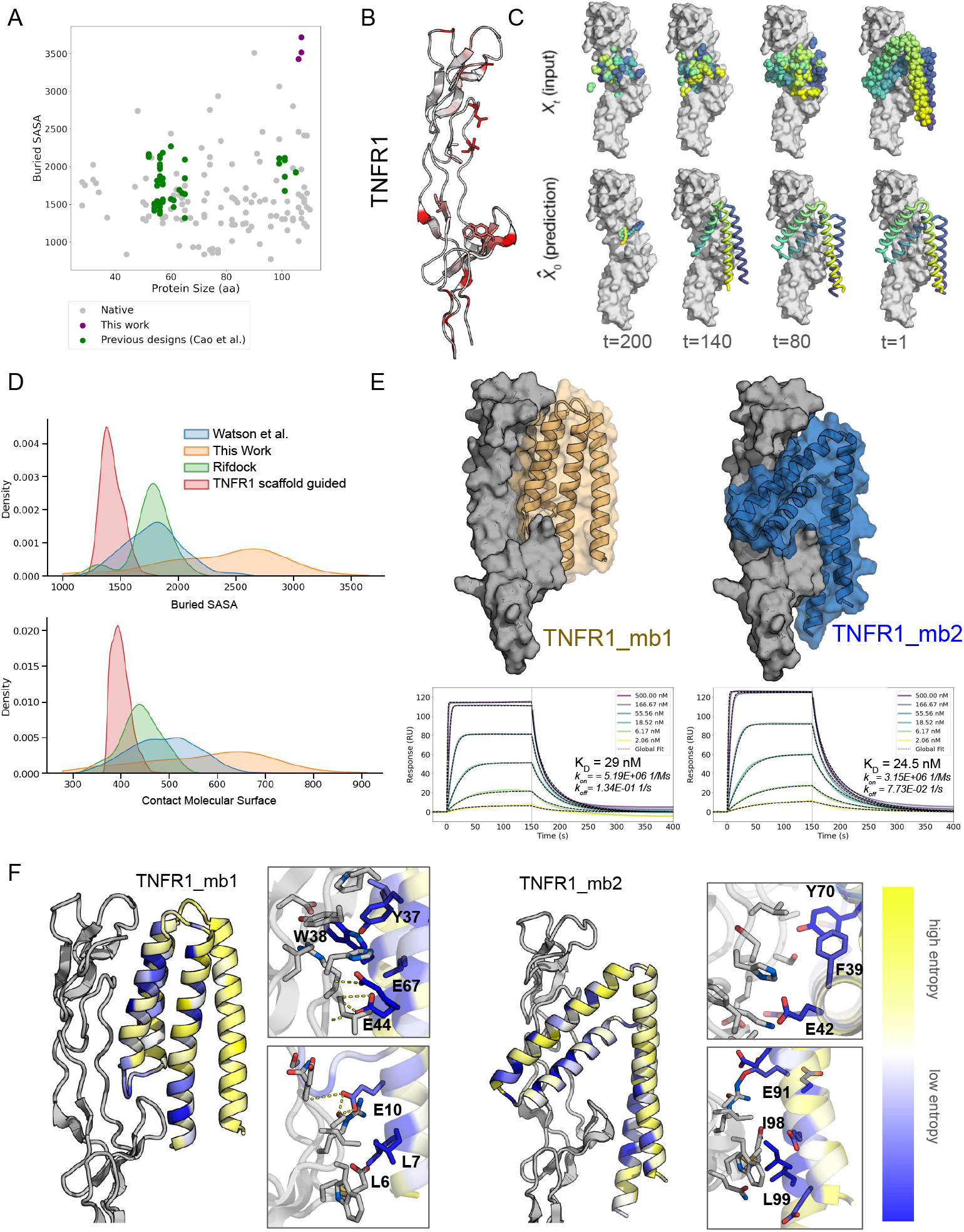
Diffusion of shape complementary binding proteins. A. Previous de novo designed binders (green) generated by docking pregenerated scaffolds bury less surface area (□^2^) against their target than many native complexes (orange); the approach developed here enables the design of very large interfaces (blue). B. TNFR1 is a challenging target, with a flat surface and few surface hydrophobic residues which are shown in red on the structure (PDB ID: 6KP8). Residues selected as target hotspots for RFdiffusion are shown as sticks. C. Representative RF_Diffusion trajectory against TNFR1 starting from a random residue distribution placed against the target (top left). At each denoising step (Xt, top), the network generates a predicted structure (^X0, bottom), and interpolates towards this structure to generate the next step (Xt-1). D. Comparison with previous TNFR1 design efforts. The RFdiffusion TNFR1 binder designs generated here (orange) have substantially higher buried SASA and contact molecular surface (CMS) than designs against TNFR1 generated previously using the Rosetta RIF dock method (red) that failed to bind. Designs generated against multiple targets using RIFdock (*3*)(green) and short chain RFdiffusion (*7*) (blue) are also shown for comparison. E. (top) Design models of binders TNFR1_mb1 (orange) and TNFR1_mb2 (blue) in complex with TNFR1 (gray). (bottom) SPR measurements of binding to TNFR1. F. Site saturation mutagenesis (SSM) results confirm design models of TNFR1 binders and associated entropy. All 2014 and 2033 single amino acid substitutions for TNFR1_mb1 and TNFR1_mb2 were expressed on yeast surface and probed using FACS with biotinylated TNFR1 followed by deep sequencing. Positions that were strongly conserved (low entropy, blue) were in the core and at the binding interface, while most surface residues away from the interface had high entropy (yellow). Entropy was calculated based on the overall change of affinity. Lower entropy (blue) is related to conserved interactions in the interface or core. Zoom-ins show dense interaction networks of low entropy residues (TNFR1 is in gray).

We reasoned that the limited shape matching achievable by scaffold dependent and/or short chain approaches could be overcome by using RFdiffusion to directly generate larger proteins starting from completely random residue distributions in the presence of the target of interest without any guidance from pre-existing scaffolds. Completely unconstrained RFdiffusion can generate a wide diversity of folds and assemblies (*7*), and proteins with folds that wrap around extended helical peptides (*8*). We set out to explore whether RFdiffusion trajectories conditioned only on the structure of a folded protein target, and not biased towards any particular scaffold or limited by the number of available residues, could generate folds with shapes matched to the target that extend over a large portion of the surface, and whether such an approach could enable the generation of high-affinity binders to targets for which previous computational design efforts had failed.

## Binder design

We chose to focus on the tumor necrosis factor receptor superfamily (TNFRSFs), which includes many important drug targets, including the TNF receptor 1 (TNFR1), which plays a key role in inflammatory disease (*9*). Like other family members, TNFR has an extended flat and largely polar surface lacking the concave sites with several hydrophobic residues that previous de novo design efforts have successfully targeted (Fig. 1B)—indeed multiple attempts to generate binders to TNFR and other family members using the approach of Cao et al.(*3*) met with little success (unpublished). We first attempted to use RFdiffusion as described in Watson et al. (*3, 7*) to generate binders to TNFR1, using guidance from 65 aa scaffold libraries or limited to chains less than 65 residues but also had little success (Fig. S1).

We next set out to adapt RFdiffusion to generate backbones with a more extended contact surface that can engage dispersed surface hydrophobic residues (Fig. 1B). Protein interactions, like protein folding, are largely driven by hydrophobic interactions, and targets with only a few surface hydrophobic residues distant in space have been particularly challenging. On TNFR, the few surface hydrophobic residues are separated by distances of up to 28 □—too small for 65 residue proteins to simultaneously engage. To overcome this limitation of previous approaches, random Gaussian residue clouds of up to 120 residues were placed at the native ligand (TNF) interface, and RFdiffusion was biased to form contacts with these dispersed hydrophobic residues. This generated a variety of unique backbones quite different from those in previous campaigns, but complementary in shape to TNFR1 (Fig. S2). ProteinMPNN was used to design sequences for these backbones in complex with TNFR1 to favor both folding to the intended structure and binding to the target. To further sample around promising designs predicted by AlphaFold2 (AF2) to form complexes (pae_interaction <20), we extracted additional backbones from preceding diffusion timesteps, as the predicted structure was already close to the final structure at around timestep 120 out of 200 (Fig. 1C). The designs predicted to most strongly bind the TNFR1 (interface pae < 7.5) and fold to the target backbone (pLDDT >85) were selected for experimental characterization. The contact molecular surface (CMS) and buried solvent accessible surface area (SASA) of these designs was substantially higher than designs from previous design campaigns (Fig. 1D).

Genes encoding 96 designs were obtained, and the proteins expressed in E coli. Despite the less regular structures and longer lengths than most previous binder designs, 90 of 96 expressed well and were primarily monomeric (Fig. S3A). Six of the designs bound to TNFR1 in surface plasmon resonance (SPR) experiments (Fig. S3B), with designs TNFR1_mb1 and TNFR1_mb2 having K_D_s of 29 nM and 24.5 nM respectively. Both designs were highly specific, with no detectable binding to TNFR2. The tertiary structure was quite distinct from previously designed *de novo* binders, and the designs interact with TNFR1 over an extended region (Fig. 1D). Design TNFR1_mb2 has an unusual V shaped fold with a very high contact molecular surface (CMS) for TNFR of 795 □^2^, substantially more than the average 490 □^2^ of previous diffused designs (*7*). Design TNFR1_mb1 also has a binding mode less regular than that of previous de novo designed binders with a connecting loop inserted in the TNFR1 binding cleft, and a similarly high CMS of 897 □^2^. For both designs, we obtained a high resolution binding footprint by determining the effect on binding of every amino acid substitution at every position one at a time (4047 substitutions in total) (Fig. S4 and S5). The results were closely consistent with the design models, with substitutions impacting binding concentrated at the designed interface and in the protein core where they would disrupt folding. The most conserved interactions for both designs centered around the hydrophobic patches involving TNFR1 residues 107/111 and residues 38/40 (Fig. 1E) spanned by more polar interactions in the center of the interface. Taken together, the close agreement of the site saturation mutagenesis (SSM) footprints with the design model interfaces, and AF2 and RoseTTAFold2 (RF2) complex predictions with the designed structures (pAE interface of 5.1 and 4.08, CA RMSD to design of 0.6 and 0.97 for designs TNFR1_mb1 and TNFR1_mb2 respectively, using AlphaFold2-multimer model 1), suggest that both TNFR1_mb1 and TNFR1_mb2 bind to TNFR1 as designed.

### In silico affinity maturation

Tumor necrosis factor (TNF-α) is a trimer that binds TNFR1 with high (19 pM) affinity (*10*), and to effectively outcompete this interaction with a monomeric protein to counter inflammation even higher binding affinities are required. To further optimize the affinity of the TNFR1_mb2 and TNFR1_mb1 designs, and the TNFR1_mb3 design, which also bound specifically (Fig. S3B), rather than combining beneficial substitutions from the SSM libraries, which requires considerable experimental screening (*3*), we used partial diffusion (Fig. 2A): backbones were partially noised (over 15 - 25 steps; 50 steps yield a completely random distribution), followed by RFdiffusion denoising which yielded new backbones resembling but distinct from the original designs (RMSD 0.58 to 4.96). We generated 25,000 partially diffused backbones around each starting structure, and following ProteinMPNN, the 32 designs for each starting structure which AF2 most confidently predicted bound TNFR1 in the designed binding mode (pae_interaction < 5) were selected for experimental characterization. These partially diffused designs had substantially greater CMS and buried SASA than the parent designs in all three cases (Fig. 2B). The designs were expressed in *E*.*coli*, and TNFR binding was measured by SPR. Most (94 out of 96) of the designs were expressed at high levels, and 30% (28 out of the 94 expressed designs) bound TNFR1 (Fig. S6A). Partial diffusion increased the binding affinity of TNFR1_mb2 by three orders of magnitude to <10 pM (Fig. 2C, S6B), while TNFR1_mb3 affinity increased from weak binding in the uM range to 20 nM (Fig. 2C). Improvements were smaller for the more regular TNFR1_mb1 backbone; free diffusion sampling appears to have already found a close-to-ideal solution for this binding mode. For TNFR1_mb2, the considerable increase in affinity brought about by partial diffusion likely is due to an additional interface and an overall better fit to the binding cleft with several additional contacts (Fig. 2C). The low picomolar affinity of the partially diffused TNFR1_mb2, which we refer to as TNFR1_mb2_pd1 below, is considerably higher than any previously described monomeric TNFR1 binding protein.

**Fig. 2.**
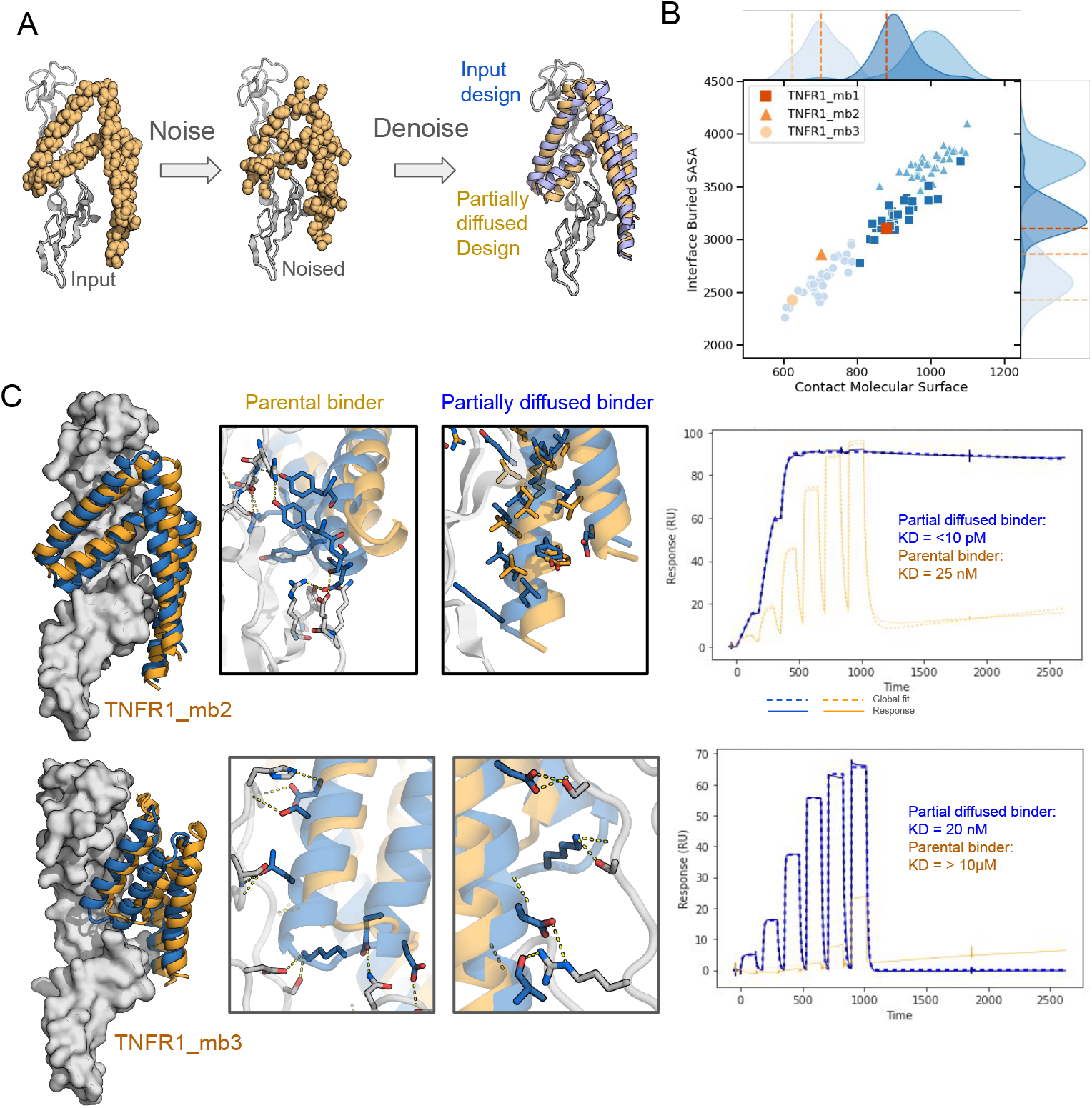
Partial diffusion generates picomolar binders. A. Schematic of partial diffusion process for TNFR1_mb2. The backbone of the input structure is represented as a collection of independent residues (first panel), noise is added (second panel), and then RFdiffusion is used to remove the noise, which results in a similar but better fitting model (orange, right) compared to the input structure (purple). B. Partial diffusion increases interface contacts. Contact molecular surface and interface buried solvent accessible surface area is depicted for input designs (orange) and the respective partial diffused variants in blue squares (TNFR1_mb1), triangles (TNFR1_mb2), and circles (TNFR1_mb3). C. Partial diffusion increases interface interaction density and binding affinity. For TNFR1_mb2 an additional interface forms (left panel), while an existing interface remains largely unchanged (right). For TNFR1_mb3, improved shape matching leads to additional interactions (bottom middle insets). The corresponding TNFR1 SPR traces are on the right (6 steps of 5x dilutions from 500 nM).

### Switching specificity

Given the success of partial diffusion in increasing binding affinities, we investigated whether a similar approach could be used to switch specificity to other TNFR family members, which are diverse in sequence, but have a very similar overall fold (Fig. 3A and B). We placed binders TNFR1_mb2, TNFR1_mb1, and TNFR1_mb3 on TNFR superfamily members TNFR2, OX40, and 4-1BB (by superimposing the latter receptors on the TNFR1 in the design models), and carried out 25,000 noising and design trajectories for each combination consisting of the addition of random Gaussian noise, RFdiffusion denoising, and ProteinMPNN sequence design. For TNFR2, 1323 designs had AF2 complex predictions with pae_interaction <7.5—a substantially higher success rate than achieved by free diffusion on TNFR1. For each of the receptors, 48 designs were experimentally characterized. For TNFR2, 32% of the designs bound with high specificity (Fig. S7A); the highest affinity design had a K_D_ of 198 pM to TNFR2 and no affinity for the other family members tested (Fig.3, Fig. S7B). Unlike TNFR2, which shares a common ligand with TNFR1 (TNF-α), OX40 has a different ligand and thus a more distinct binding interface; despite this difference in natural ligand partial diffusion starting from the TNFR1 binders yielded an OX40 binder with a K_D_ of 30 nM. As expected, this binder was shifted more substantially both in approach angle and tertiary structure compared to the parental design than the TNFR2 binders (Fig. S7C). For the even less related 4-1BB, an additional round of partial diffusion was required to achieve a pae_interaction less than 7, but the experimental success rate was still high, with 22 out of 48 tested designs binding specifically to their target, with the highest affinity of 44 nM. Design 4-1BB_mb1 illustrates how partial diffusion can conform backbones to a target, in this case, by introducing an unusual kinked helix and a short betasheet to pair with the unique receptor fold (Fig. S7F).

**Fig. 3.**
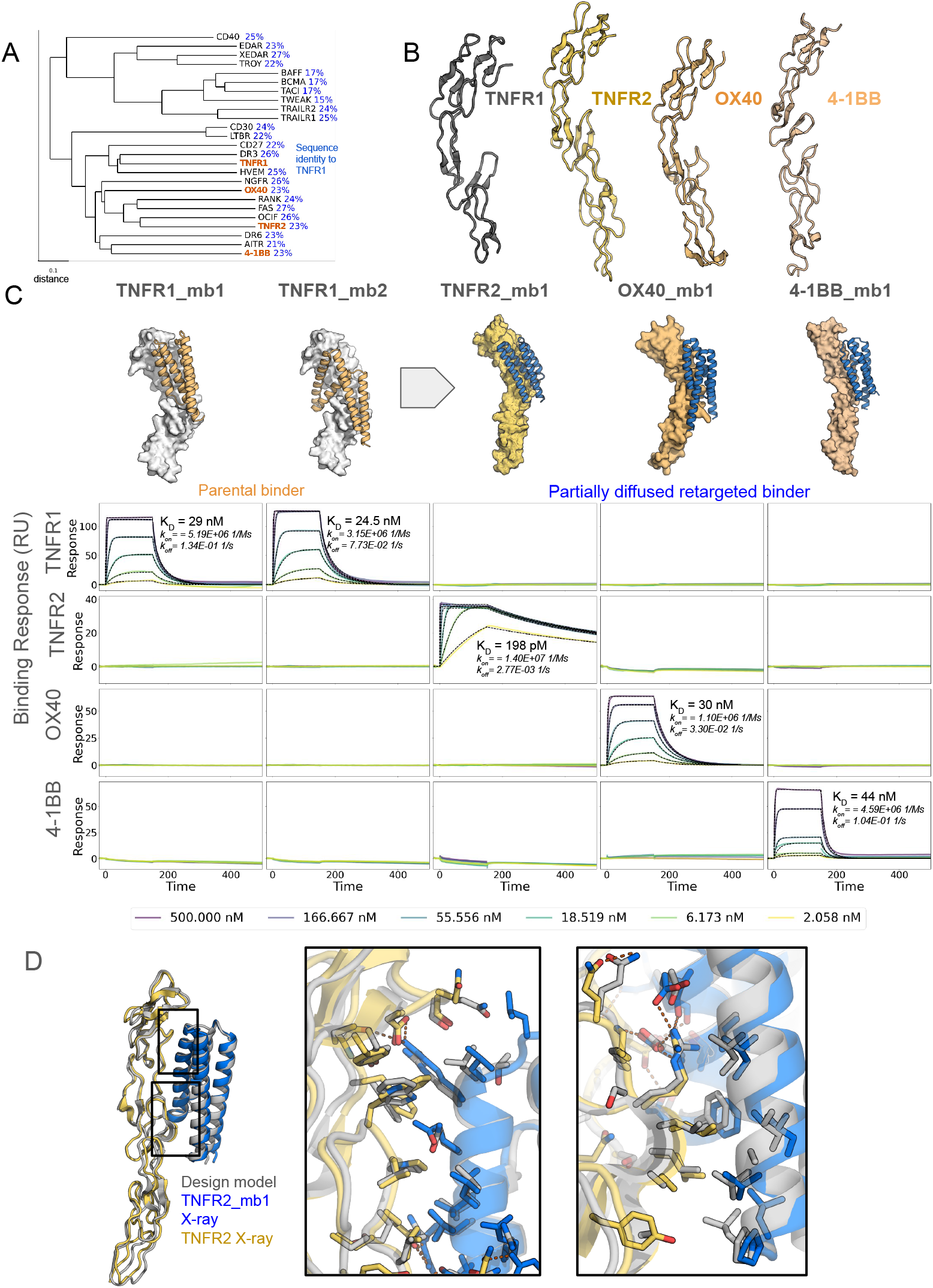
Partial diffusion generates high-specificity TNFR2, OX40, and 4-1BB binders. A. Phylogenetic tree of TNFT superfamily receptors. The tree was constructed using the UPGMA method based on a multiple sequence alignment performed with ClustalW. TNFR1, TNFR2, OX40, and 4-1BB investigated in this study are highlighted in orange. The scale bar represents a distance of 0.1 substitutions per site. Pairwise sequence identity to TNFR1 is indicated in blue. B.Comparison of structures of TNFR1, TNFR2, OX40, and 4-1BB (PDB IDs: 7KP8 (*17*), 3ALQ (*17, 18*), 2HEV (*19*), 6BWV (*20*)). C. (Top) Original design models and partial diffusion generated model for the highest affinity TNFR2, OX40 and 4-1BB binders (also see Fig. S6). The two TNFR1 binders (orange, left) were superimposed on the new targets (right) and partially diffused to yield target-matched backbones (blue). (Bottom) SPR measurements show that the binders are highly specific for the targets they were diffused against. D. Crystal structure (colors) of TNFR2_mb1 (blue) in complex with TNFR2 (yellow) superimposed on the computationally design model (grey). Boxed interface regions on the backbone superposition on the left are shown with interface sidechains in the zoom ins on the right.

### TNFR2 design has near atomic level accuracy

We were able to solve the structure of the highest affinity retargeted binder, TNFR2_mb1, in complex with TNFR2 using X-ray crystallography. The crystal structure of the complex is very close to the computational design model, and nearly identical over the designed binder (C-alpha RMSD 0.52□; Figure 3D, left panel). The key sidechain interactions in the extensive designed protein protein interface are very similarly positioned as well (Figure 3D, right panels).

### Designed binders antagonize signalling

The picomolar affinity of the TNFR1 binders makes them possible candidates for blocking inflammation. To date, targeting of the TNF-α pathway has primarily focused on binding to circulating TNF-α due to the fact that antibodies targeting TNFR1, owing to their bivalency, can activate, rather than suppress, TNF-α signaling. We investigated if our (monomeric) designs could inhibit TNF-α signaling on a TNF-α HEK293 reporter cell line (InvivoGen), which monitors TNF-α signaling via AP-1/NF-κB dependent activation of a SEAP reporter gene (Fig. 4A). We found that the designs potently inhibited TNF-α signaling, with an IC50 for the best design at 106 pM.

**Fig. 4.**
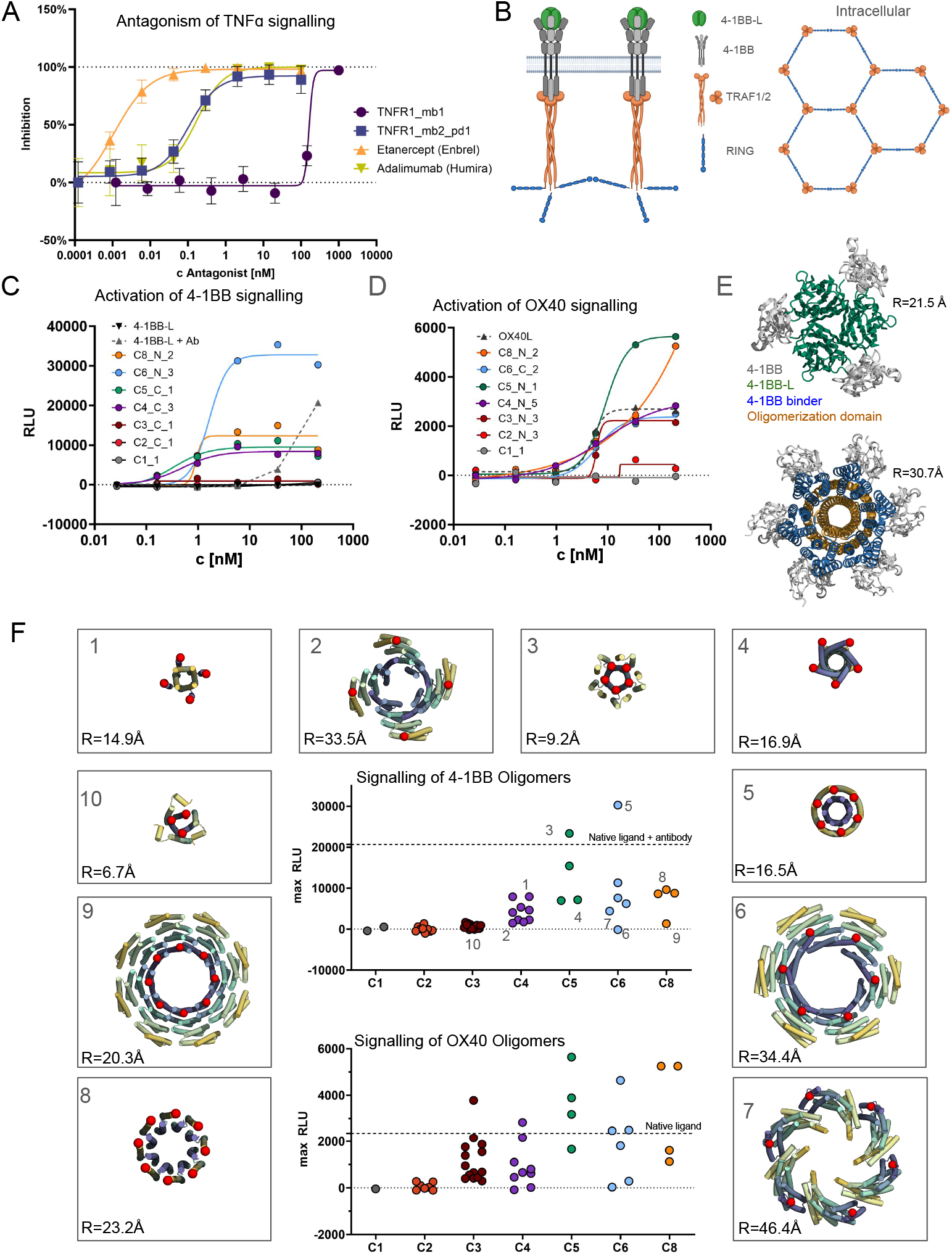
Design of soluble oligomeric 4-1BB and OX40 superagonists. A. Designed binders antagonize TNF□ signaling. HEK293-blue cells were incubated with 100 pM TNF-α and serial dilutions of designed binders and NFkB-dependent activation were measured. Curves are fit to data from two independent replicates. B. Schematic overview of 4-1BB signaling. Clustering of trimeric 4-1BB Ligand (4-1BB-L, green) with three copies of 4-1BB receptor (gray) leads to intracellular formation of TRAF1/2 trimers (orange) and intracellular hexamers of zinc-RING finger domains leading to downstream signaling. C. Multivalent presentation of 4-1BB binder design on cyclic oligomers activates 4-1BB signaling on luciferase reporter cell lines, compared to native 4-1BB ligand (4-1BB-L) alone and complexed with an anti-his antibody. 41BB_mb2 was fused to various oligomerization domains, examples for different cyclic oligomeric states with a valency of one for binder alone (C1_1) to valency of eight (C8_N2) are shown. For details on oligomers see also Table S4 and Figure S10. D. Multivalent presentation of OX40 binder design on cyclic oligomers activates OX40 signaling on luciferase reporter cell lines. OX40_mb1 was fused to various oligomerization domains, examples for different cyclic oligomeric states with a valency of one for binder alone (C1_1) to valency of eight (C8_N2). For details on oligomers see also Table S4 and Figure S10. E. Model of 4-1BB (gray) upon binding of 4-1BB_mb_1 (blue) fused to the N-terminus of a cyclic hexamer (orange) compared to native complex with 4-1BB-L (green) (right, PDB ID: 6BWV). Distance between receptor (M101) to center is indicated (R). F. Geometry and oligomerization state dependence of designed 4-1BB and OX40 agonists. Binders 41BB_mb2 and OX40_mb1 were fused N- or C-terminal to 24 oligomerization domains each (see also Table 2). Central panel shows maximum recorded signal at 200 pM, with each dot representing one oligomer-binder fusion, separated in groups depending on oligomeric state for C2-C8 oligomers. Each Dotted line indicates signal of the native ligand (OX40) or native ligand plus antibody (4-1BB). Examples of design models of oligomerization domains that show high or low signal in specific groups for 4-1BB are shown in the surrounding panels, with numbers indicating their signal in the central plot. Fusion sites of binders are indicated by a red sphere and distance to center (R) labeled. Chains are colored in a gradient from N-terminus (yellow) to C-terminus (blue).

Further biophysical characterization showed that the designs have desirable developability features. They retain function after heating to 95°C for 30 minutes or incubation in mouse serum for two hours (Fig. S8A, Fig. S9). While the TNFR1 binders have substantial hydrophobic patches in their target binding surfaces (see Fig 1F), hydrophobic interaction chromatography (HIC) did not show substantial hydrophobic interaction compared to Adalimumab and Etanercept and other clinical antibodies (Fig S8B). In addition to having high specificity within the family (Fig 3C), the highest affinity TNFR1_mb2_pd1 also showed no substantial off-target binding to a TNFR1 knockout cell line up to 100 nM, 10,000 fold higher than the K_D_ (Fig S8C).

#### Agonist design

For OX40 and 4-1BB, which have been widely studied for expanding T cells for cancer treatment, agonists could have therapeutic potential. Unlike TNF□/TNFR1, 4-1BB signaling is not activated by soluble trimeric ligands alone–the physiological ligands are in the plasma membrane of adjacent cells, and induce 4-1BB arrangements with longer range order (Fig. 4B). Signaling has been achieved using antibody-generated ligand networks to drive higher-order complexes (*11–13*), but while such ligands are soluble, they are also quite heterogeneous. To explore the generation of well-defined monodisperse 4-1BB agonists, we fused the 4-1BB binder to designed homo-oligomers with different valencies and spacings between fusion sites (red dots in Fig. 4). We found that monomers and C2 or C3 oligomers did not signal, consistent with the lack of signaling of native trimeric ligands. In contrast, signaling was observed with C4, C5, C6, and C8 oligomers (Fig. 4C, Fig. S10A). The strongest signal was observed for a C6 construct which arranges the receptors at similar distances as the native ligand, but with higher valency (Fig. 4E). Over a set of 44 oligomers (Fig. 4F), valency was the strongest determinant of signal strength, with no signal for C1-C3, a consistent but low signal for C4, and a higher signal for C5 and C6. Beyond valency, the geometry of association also influenced the extent of signaling; higher order oligomers that separate the receptors by distances significantly greater than the native ligand exhibited low or no signal as did those that would likely clash with the receptor or penetrate the membrane (Fig. 4F). Overall, the requirement for higher order valencies is consistent with the proposed signaling mechanism inferred from the intracellular hexameric arrangement (*14*) (Fig. 4B), but it remains to be determined how bringing in just one more subunit (in the C4 case) leads to agonism.

For OX40, we again tested a range of different oligomeric states, and observed a quite different pattern. In contrast to 4-1BB, trimeric constructs were effective agonists, consistent with the fact that OX40 can be activated by soluble trimeric ligands. Monomeric and dimeric binder constructs did not signal, while oligomeric constructs with three, four, or five binding modules efficiently activated signaling (Fig. 4D, F, Fig. S10B).

For both 4-1BB and OX40, both the EC50 and Emax (maximum signal) varied considerably among the oligomeric constructs, indicating a substantial opportunity for fine-tuning the response by modulating valency and geometry, both to investigate the mechanism of signaling through this important class of receptors, and for therapeutic applications. Particularly interesting are the substantially higher Emax of the best OX40 and 4-1BB synthetic agonists compared to the native ligand in the OX40 case and antibody-ligand assemblies in the 4-1BB case; these could be particularly useful for expanding T-cell populations.

## Discussion

For therapeutic challenges for which antagonism without any risk of agonism is necessary, high-affinity monomeric binders could have advantages over bivalent antibodies, which can potentially dimerize the target receptor and activate signaling (*9, 15*). TNF-α and TNFR are key drug targets given the central role this interaction plays in inflammatory disease; current therapies primarily target the ligand TNF-α instead of TNFR1 to avoid potential activation of inflammatory responses, but binding TNF-α also inhibits potentially anti-inflammatory signaling through TNFR2, which could contribute to unwanted side effects of this important class of drugs (*16*). The very large interfaces of the TNFR1 binders, even larger than the native interface despite being only 107 amino acids, could likely enable even higher affinity antagonism into the fM range, and the high stability and likely low cost of production of small designed proteins could enable oral administration for gut disease. As illustrated by our 4-1BB and OX40 superagonists, the high affinity monomeric binders enable the construction of a wide variety of soluble signaling molecules, offering far more control than current native ligand plus crosslinking antibody-based approaches.

More generally, the ability to generate high affinity and specificity binders to therapeutically important and structurally challenging protein targets without having to immunize animals, screen large random libraries, or test thousands of design candidates ushers in a new era for binder design and therapeutic candidate discovery. The number of sequences tested (96 free diffusion and 96 partial diffusion designs for TNFR1, and 48 partial diffusion designs for TNFR2, OX40, and 4-1BB) is far fewer than in previous studies in which libraries of tens of thousands of designs were screened using yeast display, and no random or experimentally guided optimization was involved other than selecting the best of the 96 first round designs for partial diffusion. The <10 pM affinity for TNFR1 and 198 pM for TNFR2 are the highest we are aware of for monomeric binders to these targets; for comparison, the antibodies that likely bind bivalently have affinities of up to 680 pM, while monomeric Fabs, scFvs, and nanobodies are in the range of 10-100 nM (*15*). This high affinity for targets for which multiple previous binder design efforts failed likely reflects the very high designed shape complementarity and buried surface area. Indeed, the amount of surface area these designs bury on their targets is substantially higher than in previous minibinder design efforts, and the buried surface area per residue rivals that of the native complexes, which have evolved over hundreds of millions of years (Fig. 1A). The combination of RFdiffusion starting from random residue clouds of >100 amino acids and partial diffusion to optimize affinity and achieve high specificity for family members provides a very powerful and general approach to obtaining high potency and affinity binders to challenging classes of targets.

## Acknowledgments

We thank S. Berger, J. Lazarovitz, G. Ueda and I. Swanson Pultz for helpful discussions; K. VanWormer and L. Goldschmidt for technical support. We thank M. Kennedy for reading and editing the manuscript. We thank B. Wicky and L. Milles for support in high-throughput protein expression and for generation of oligomerization domains. Crystallographic work is done at the National Synchrotron Light Source 2 on beamline FMX (17-2). We thank Amgen for kindly gifting recombinant TNFR2 for crystallization.

## Author Contributions

D.B. directed the work. M.G. designed, screened and experimentally characterized all binders. A.Kr. experimentally characterized signalling of binders. R.J.R. developed the oligomer screen and performed and advised on experiments. I.G. and M.G.S. performed yeast display screening. B.C. developed computational tools and analyzed binders. A.K.B., A.Ka. and E.J. crystallized complex and solved the TNFR2_mb1 structure. G.A. generated knock-out cell lines and conducted cell surface staining. B.H. and W.Y. advised on binder design and designed site saturation mutagenesis libraries. W.C. advised on and performed antagonism experiments. B.K. supported design of 4-1BB binders. M.G. and D.B. wrote the manuscript, which all authors reviewed and commented on.

## Data availability

All data are available in the manuscript or the supplementary materials. Atomic coordinates, and structure factors reported in this paper have been deposited in the Protein Data Bank (PDB), http://www.rcsb.org/ with accession code **9CU8**.

## Supplementary Materials

Materials and Methods

Figs. S1 to S10

Tables 1-4

