## Supplemental Material for "Target-conditioned diffusion generates potent TNFR superfamily antagonists and agonists"

**Supplementary Materials**
Materials and Methods
Figs. S1 to S10
Table 1 to 4

### Materials and Methods

#### Design of protein binders to TNFRSFs

To design de novo binders to TNFRSFs using RFdiffusion, we initially generated backbones to TNFR1 based on a cryo-EM structure (PDB ID: 7KP8). We generated 30,000 initial designs using RFdiffusion(*7*). We targeted binders using input "hotspot" residues to a specific site on the target protein. The hotspots selected were as follows (chain and residue index from PDB): F67, F70, F95, F112.

In line with current best practice, we used the ProteinMPNN-FastRelax protocol described in Bennett et al (*21*), this protocol starts with a round of ProteinMPNN and then cycles between FastRelax and ProteinMPNN to attempt to iteratively improve the sequence and structure agreement. Two sequences per design were generated. To filter designs we ran AF2 with an initial guess and target templating. Briefly, this configuration of AF2 runs without a multiple sequence alignment and without template information for the de novo binder, which ensures that predictions are not biased towards examples that have sequence or structural homology to the PDB. This configuration uses the template feature in AF2 to provide the exact structure of the target protein, as we are designing with a rigid target and know the structure of the target a priori, we desire for AF2 to keep the target fixed and only predict the dock and structure of the de novo binder. Finally, this configuration of AF2 initializes the dock and structure of the de novo binder with the RFdiffusion design model of the dock and structure.

For designs that were predicted as complex with pae_interaction <20, multiple backbones were extracted from the trajectories (10 backbones at every 5$^{th}$ timestep from 0-50), and for the resulting 1210 backbones the ProteinMPNN-FastRelax-AF2 pipeline repeated, to diversify designs and achieve a sufficient number to order on a 96 scale.

*Partial diffusion*

RFdiffusion was modified to allow the input structure to be noised only up to a user-specified time step instead of completing the full noising schedule as described previously (*8*). The starting point of the denoising trajectory is therefore not a random distribution, but contains information about the input distribution resulting in denoised structures that are structurally similar to the input. The AF2 models of the initial binders TNFR1_mb1-3 were used as inputs to partial diffusion, either directly or after superimposing and replacing the receptor with TNFR2,

OX40, or 4-1BB (PDB IDs: 3ALQ, 2HEV, 6BWV respectively). The models were subjected to 15 or 25 noising time steps out of a total of 50 time steps in the noising schedule, and subsequently denoised. An auxiliary potential to promote interface contacts was used in some cases. This interface potential artificially pushes the coordinates of the binder closer to the target according to the gradient of a heuristic function over the whole trajectory, and is described in detail in the public RFdiffusion repository under https://github.com/RosettaCommons/RFdiffusion/blob/main/rfdiffusion/potentials/potentials.py The decay (linear or quadratic) of this potential was variable for subsets of the design campaigns. Approximately 20,000 partially diffused designs were generated for each target. The backbones in the resulting library were sequence designed as described above for free diffusion designs. The designs with the lowest pAE_interaction and pLDDT >85 were selected for experimental characterization.

**Plasmid construction**

Protein binder designs were ordered as synthetic genes (eBlocks, Integrated DNA Technologies) with compatible BsaI overhangs to the target cloning vector, LM0627 (*22*) for Golden Gate assembly. LM0627 is a modified expression vector containing a Kanamycin resistance gene and a ccdb lethal gene between BsaI cut sites. Subcloning into LM0627 results in the following product: MSG-[protein]- GSGSHHWGSTHHHHHH, with the C-terminal SNAC cleavage tag and 6XHis affinity tag respectively underlined.

Oligomerized binders were generated by cloning those eBlocks in similar vectors which resulted in both C- and N- terminal fusion to 24 oligomerization domains, and in a final construct of MSG-[binder]-GSGS-[oligomerization domain]-GSGSHHWGSTHHHHHH or [oligomerization domain]-GSGS-[binder]-GSGSHHWGSTHHHHHH. Oligomerization domains were described previously (*22*, *23*), a full list of constructs can be found in Table 3.

**Protein expression and purification**

For protein binder expression screens, a previously reported protocol (*22*) was followed with some modifications as denoted. In short, Golden Gate subcloning reactions of designs were carried out in 96-well PCR plates in 1 µL volume. Reaction mixtures were then transformed into a chemically competent expression strain (BL21(DE3)), and 1-hour outgrowths were split directly into four 96-deep well plates containing 1 mL of auto-induction media (autoclaved TBII media supplemented with Kanamycin, 2 mM $MgSO_4$, 1X 5052) for a final total volume of approximately 4 mL. The following day (20-24 hrs later), cells were harvested and lysed, and clarified lysates were applied directly to a 100 µL bed of Ni-NTA agarose resin in a 96-well fritted plate equilibrated with a Tris wash buffer. After sample application and flow through, the resin was thoroughly washed, and samples were eluted in 200 µL of a Tris elution buffer containing 500 mM imidazole.

All eluates were sterile-filtered with a 96-well 0.22 µm filter plate (Agilent 203940-100) prior to size exclusion chromatography (SEC). Protein designs were then screened via SEC using an

AKTA FPLC outfitted with an autosampler capable of running samples from a 96-well source plate. Protein binders were run on a Superdex75 Increase 5/150 GL column (Cytiva 29148722), and symmetric oligomers were run on a Superdex200 Increase 5/150 GL column (Cytiva 28990945). For all binding proteins, HBS-EP+ ( 0.01 M HEPES pH 7.4, 0.15 M NaCl, 3 mM EDTA, 0.005% v/v Surfactant P20) was used as a running buffer for subsequent SPR analysis without buffer exchange. For oligomerized binders, 20 mM NaPhos pH 7.4, and 100 mM NaCl was used as a running buffer, 0.25 mL fractions were collected from each run, and selected fractions were pooled for further analysis.

For larger scale protein purification, protein expression was performed using 50 mL of auto-induction media (autoclaved TBII media supplemented with Kanamycin, 2mM $MgSO_4$, 1X 5052), and grown for 24h at 37°C. The cells were harvested by spinning at 4,000 x g for 10 min and then resuspended in lysis buffer (100 mM Tris-HCl, 200 mM NaCl, 50 mM imidazole). Then, the cells were lysed by sonication in a Qsonica, Q500 with a 4-pronged horn for 2:30 min ON total, with an amplitude of 80%. Soluble fractions were clarified by centrifugation at 4,000 x g for 30 minutes, and were subsequently purified by affinity chromatography using bed Ni-NTA resin (Qiagen or Thermo Fisher) on a vacuum manifold. A series of 3 washes using wash buffer (20 mM Tris-HCl, 200 mM NaCl, 50 mM imidazole) was performed prior to elution with elution buffer (20 mM Tris-HCl, 200 mM NaCl, 500 mM imidazole). After elution, protein samples were filtered and injected into an autosampler-equipped Akta pure system on a Superdex S75 Increase 10/300 GL column(Cytiva 28-9909-44) at room temperature. The SEC running buffer was 20mM Tris-HCl, 100mM NaCl pH 8.

**Crystallography sample preparation and data collection**

For crystallization TNFR2_mb1 was expressed and purified as described above. Following affinity purification, size exclusion chromatography (SEC) was performed in SNAC cleavage buffer (100 mM CHES, 100 mM NaCl, 100 mM acetone oxime, 500 mM guanidine HCl, pH 8.6). 2 mM of $NiCl_2$ was added and the solution was incubated overnight at 37°C. Following cleavage, the solutions containing the cleaved protein products were incubated with 1 mL Ni-NTA resin to bind any uncleaved product, and the flow through was Collected. Cleaved TNFR2_mb1 was complexed at twofold molar excess with TNFR2 ectodomain, and the complex purified using size exclusion chromatography (SEC) using 20 nM Tris/HCl 100 nM NaCl at pH 8 as running buffer. Sample was then concentrated to 13 mg/ml, and subjected to hanging drop vapor diffusion crystal screen at a drop size of 200 nl and Buffer/Sample ratios of 1:2, 1:1 and 2:1.

Crystals grew successfully in 1.0M Imidazole; MES monohydrate (acid) pH 6.5, 0.12 M Monosaccharides, 40% v/v PEG 500 MME; 20% w/v PEG 20000. Crystals were harvested directly from a screening tray, and flash cooled in liquid nitrogen. X-ray diffraction was performed at NSLS2 beamline 17-2, data were processed with XDS (*24*), and merged/scaled using Pointless/Aimless in the CCP4 program suite (*24*, *25*). The structure was phased by molecular replacement using the designed structure as the search model by Phaser (*26*) and refined with Phenix. Following molecular replacement, the models were improved, and efforts were made to reduce model bias. Structures were refined in Phenix (*27*). Model building was performed using COOT (*28*). The final model was evaluated using MolProbity (*29*). Data collection and refinement statistics are recorded in Table S3. Data deposition, atomic coordinates, and structure factors reported in this paper have been deposited in the Protein Data Bank (PDB), http://www.rcsb.org/ with accession code **9CU8**.

**Surface Plasmon Resonance**

Binding kinetics were analyzed via Surface Plasmon Resonance (SPR) on a Biacore 8K (Cytiva). Binding to various TNFRSFs was measured by capturing Fc-tagged receptor ectodomain of TNFR1, TNFR2, OX40, and 4-1BB (all Sino Biological, #10872-H02H-100, #10417-H03H, #10481-H02H, #10041-H02H) or biotinylated receptor (all Sino Biological, #10481-H49H-B, #10417-H27H-B, #10417-H27H-B, #10872-H27H-B) on a Protein A chip (Cytiva #29127556) or captured Streptadvidin using Biotin CAPture Kit (Cytiva #28920234) respectively, by injecting 0.125 µg/mL receptor at a flow rate of 10 µL/min in HBS-EP+ (0.01 M HEPES pH 7.4, 0.15 M NaCl, 3 mM EDTA, 0.005% v/v Surfactant P20, Cytiva #BR100669) aiming for a capture level of ~250 response units. Analytes were diluted in HBS-EP+ and injected at a flow rate of 30 µL/min to monitor association. HBS-EP+ was used as a running buffer during dissociation at a flow of 30 µL/min. Screens were performed at a single dilution, full kinetics by running single cycle kinetics, injecting increasing concentrations of ligand or parallel kinetics using different concentrations in the eight channels. Association/dissociation times and concentration ranges were varied to suit the respective analytes. Binding kinetics were determined by global fitting of curves to $k_{on}$ and $k_{off}$ assuming a 1:1 Langmuir interaction, using the Cytiva evaluation software.

**Yeast surface display**

Saccharomyces cerevisiae EBY100 strain cultures were grown in C-Trp-Ura medium supplemented with 2% (w/v) glucose. For induction of expression, yeast cells were centrifuged at 6,000g for 1 min and resuspended in SGCAA medium supplemented with 0.2% (w/v) glucose at the cell density of $1 \times 10^7$ cells per mL and induced at 30 °C for 16–24 h. Cells were washed with PBSF (PBS with 1% (w/v) BSA) and labeled with biotinylated TNFR1 (R&D Systems #AVI10910). The cells were first incubated with biotinylated targets, washed and secondarily labeled with anti-c-Myc fluorescein isothiocyanate (FITC, Miltenyi Biotech) and streptavidin–phycoerythrin (SAPE, ThermoFisher). For SSM libraries, two rounds of sorts were applied and in the third round of screening, the libraries were titrated with a series of decreasing concentrations of targets to enrich mutants with beneficial mutations.

**Analysis of site saturation mutagenesis data and entropy calculation**

NGS data and apparent yeast SC50 was analyzed with script as described in detail in Cao et al. 2022. Briefly, raw counts were transformed into apparent yeast KD/SC50 (sorting concentration 50), the script outputs the upper and lower bounds for the SC50 (sorting concentration 50) where 50% of binder variants from that design would be collected. Lower bound KDs were reported in this study.

For each position on the binder, the sequence entropy (Shannon entropy) of each position was calculated using the observed frequencies of each amino acid in the NGS. The specific pool that was chosen for this analysis was the pool with concentration closest to tenfold lower than the calculated SC50 of the parent.

The scripts used for SC50 calculation from NGS data and entropy calculator based on SC50 was published previously in Cao et al. and is accessible under https://files.ipd.uw.edu/pub/robust_de_novo_design_minibinders_2021/supplemental_files/scripts_and_main_pdbs.tar.gz

**Hydrophobic interaction chromatography**

Minibinders and clinical proteins were buffer exchanged into 1 M ammonium sulfate and 0.1 M sodium phosphate at pH 6.5. A salt gradient was established on a ProPac™ HIC-10 (Thermo Fisher #063655) from 1.8 M ammonium sulfate, 0.1 M sodium phosphate at pH 6.5 to the same condition without ammonium sulfate. The gradient ran for 17 min at a flow rate of 1 ml/min. An acetonitrile wash step was added at the end of the run to remove any remaining protein and the column was re-equilibrated over 7 column volumes before the next injection cycle. Peak retention times were monitored at A280 absorbance.

**Circular Dichromatism**

Far-ultraviolet circular dichroism measurements were carried out with a JASCO-1500 instrument equipped with a temperature-controlled multi-cell holder. Wavelength scans were measured from 260 to 190 nm at 15 °C steps from 25 to 95 °C and back to 25 °C after refolding. Temperature melts monitored the dichroism signal at 222 nm in steps of 2 °C min–1 with 30 s of equilibration time. Wavelength scans and temperature melts were performed using 0.2 mg/ml protein in PBS buffer (20 mM $NaPO_4$, 150 mM NaCl, pH 7.4) with a 1 mm path-length cuvette

**Generation of TNFR1 KO cells**

HeLa cells were cultured in Dulbecco's minimum essential medium (DMEM) supplemented with 10% heat-inactivated fetal bovine serum (HI-FBS) and 1% penicillin/streptomycin. To generate TNFR1 knockout (KO), wild-type (WT) HeLa cells were nucleofected with a Lonza 4D-Nucleofector Unit using the Lonza SE Cell Line 4D-Nucleofector Kit. sgRNAs and spCas9 were obtained from Synthego's Gene Knockout Kit v2, and nucleofection was carried out using manufacturer's protocol.

**Cell-surface binding in WT and TNFR1 KO cells**

WT and KO cells were trypsinized and transferred to 96-U bottom plates. The cells were washed twice with ice-cold FACS buffer (0.02% BSA, DPBS) and incubated on ice with varying concentrations of TNFR1 minibinder for 30 minutes. The cells were then washed once with FACS buffer, and stained with anti-His-antibody-647 on ice for 30 minutes. The cells were washed twice, stained with Sytox Green, and analyzed on the Attune flow cytometer (Thermo Fischer).

#### TNF-α signaling assays

HEK-Blue-TNF-α cells (Invivogen #hkb-tnfdmyd) were cultured in DMEM media and maintained according to standard protocol. On Day 0, cells were plated at a density of 50,000 cells per well in a Corning 96 well flat bottom TC treated plate. On Day 1, ligands pre incubated with 0.01 nM TNF-a were added to the cells with concentrations ranging from 1uM to 0.008 uM. Plates were then incubated at 37°C for 24 hours to allow for TNF-α-induced activation of signaling pathways. Following incubation, the cell supernatant was collected, and 10 µL of the supernatant was mixed with 90 µL of QuantiBlue reagent (Invivogen #rep-qbs3). The mixture was incubated at 37°C for 30 minutes to allow for the conversion of the reagent, and the optical density at 615 nm (OD615) was measured using a Neo2 plate reader.

#### OX40/4-1BB signaling assays

For OX40 and 4-1BB signaling assays, commercially reporter Jurkat cells assay kits were used. (Promega #JA2191 and #JA2351) In both cases, cells were thawed, and 20,000 cells were distributed in 25 µL RPMI media supplemented with 10% FCS to 384-well plates. An equal amount of diluted binder oligomers was added to cells in RPMI media supplemented with 10% FCS. After 8h of incubation at 37°C, cells were lysed with lysis buffer supplemented with BrightGlo substrate as supplied by the manufacturer, and transferred to black 384-well plates. After 10 min of incubation, luminescence was read out with a Neo2 plate reader.

**Supplementary Figures**

Figure S1

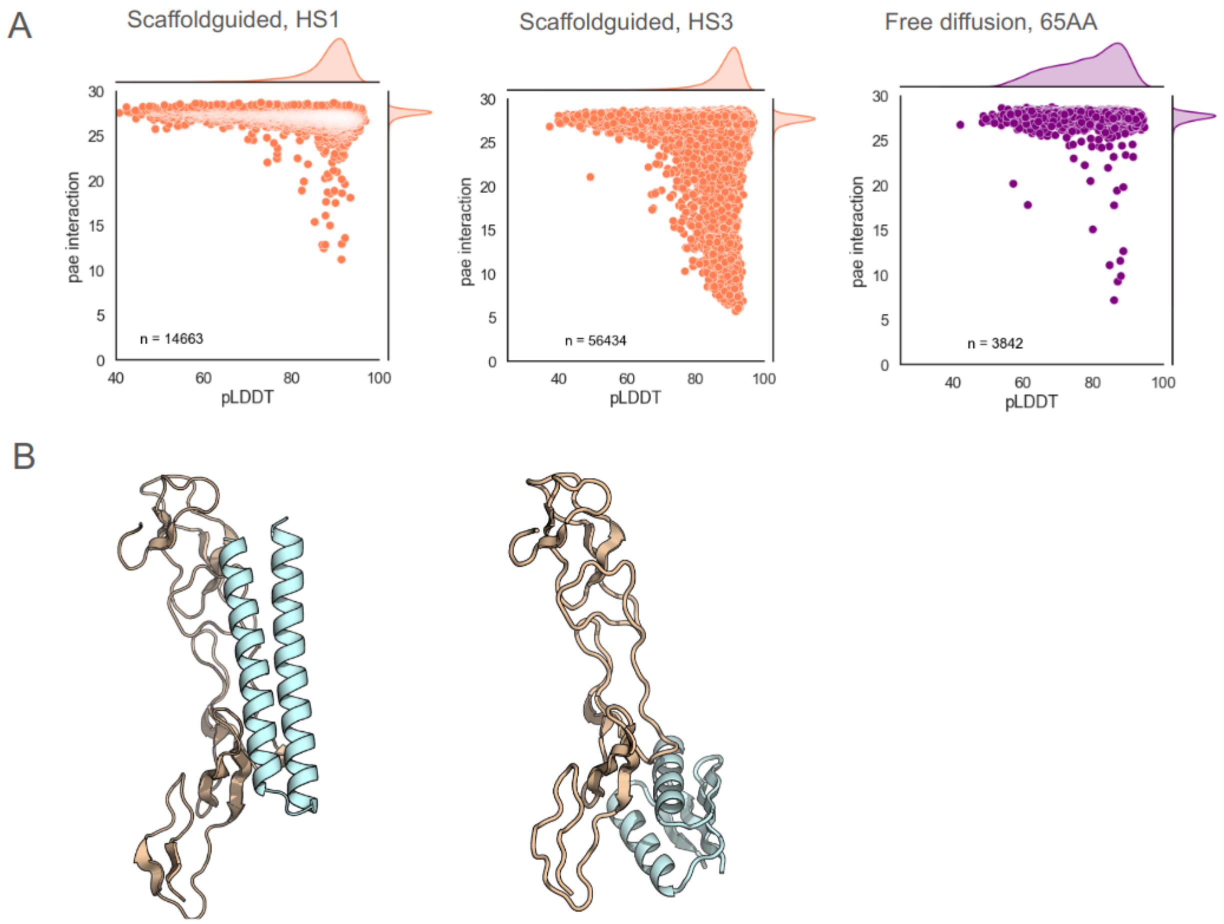


**Supplementary Fig. 1**. Scaffold guided and free diffusion of short binders fails to generate successful designs. A. Scaffold guided diffusion, in which noise is preconditioned based on a starting scaffold (described in Watson et al.) to one set of hotspot residues fails to generate designs with good predicted interaction (left), which means the pae interaction values do not reach below a threshold of 10 and are unlikely to work experimentally. A second set of hotspot residues gives better metrics, but designs are still considered a failure, since they do not respect the hotspot and move away from the native interface (see B). Free diffusion without scaffold guidance (right) fails to generate successful designs, the few designs with low pae interaction are two helical bundles which are not considered promising backbones for binder design (see B). B. Failure modes of short design diffusion. Both for scaffold guided design or free diffusion, limiting the designs to 65 residues did not results in promising designs. Examples for both failure modes are shown. Designs with good metrics either devolved to non-ideal backbones (1 or 2 helices, left) or moved away from the originally targeted interface (right).

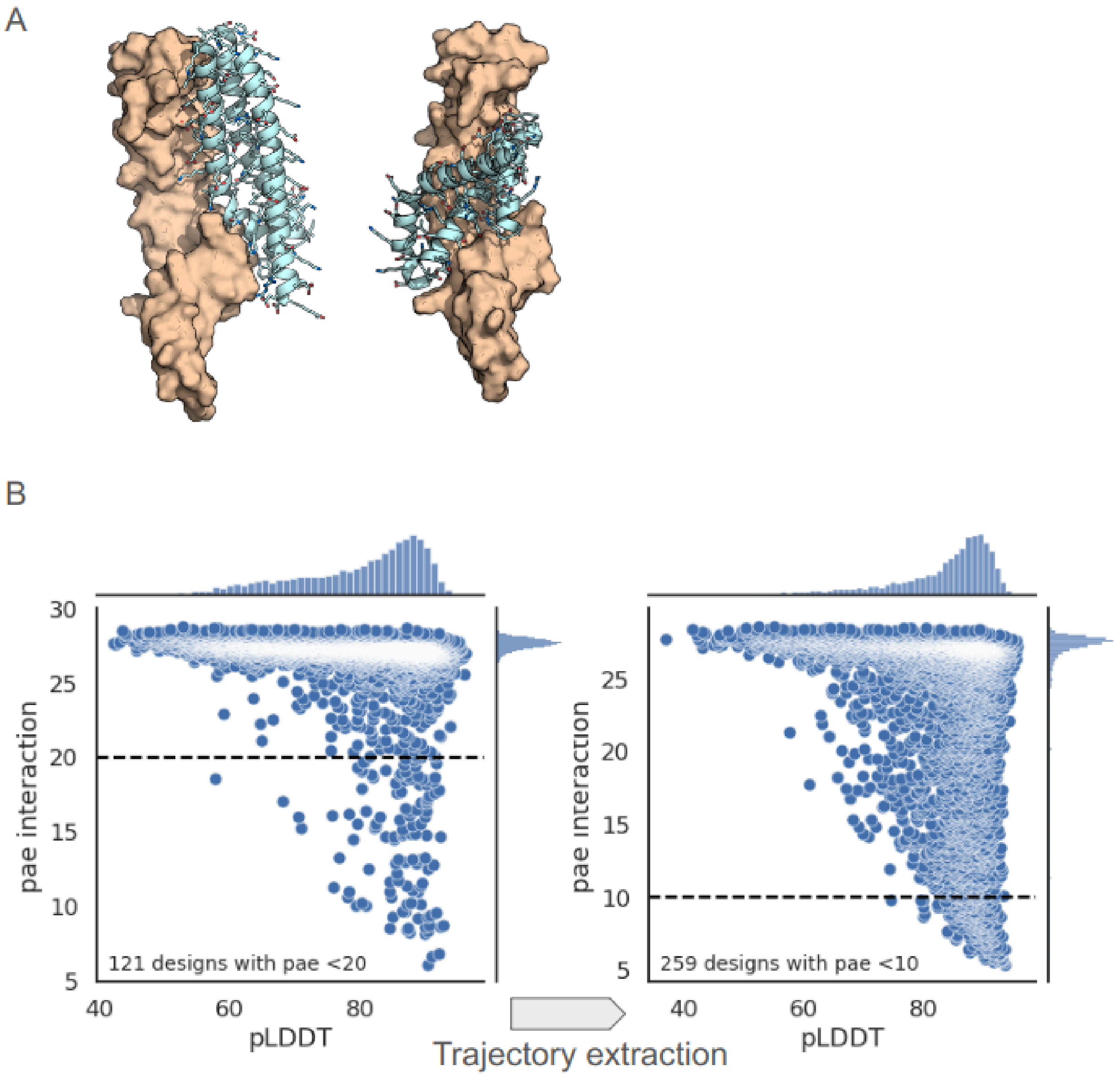


**Supplementary Fig. 2.** Free diffusion of extended binders generates promising backbones designs. A. Examples of backbones with unique folds to shape match the TNFR1 fold. B. Metrics of free diffused designs after resampling of promising backbones extracted from trajectories of designs with a pae_interaction <20.

Figure S3

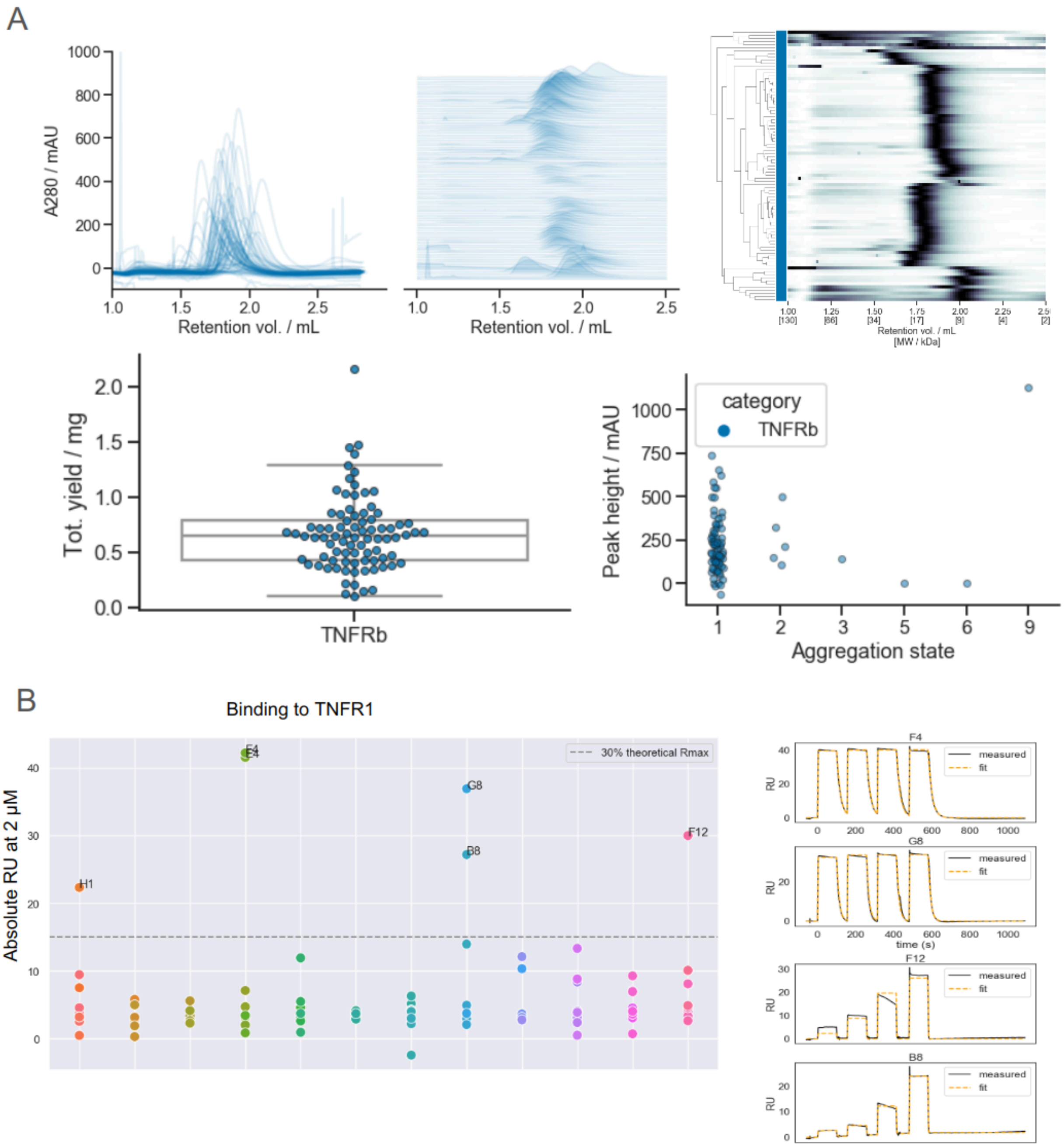


**Supplementary Fig. 3.** Expression and analysis of TNFR1 binder campaign. A. 96 TNFR1 binders were expressed from *e.coli* and purified by size exclusion chromatography (SEC). The majority was monodispersed on SEC (upper panels) and expressed to high yields without aggregation (lower panels) B. Surface Plasmon Resonance (SPR) screening of expressed binders. All binders were tested at 4 concentrations. The left panel shows the absolute response at 2 µM binder, single cycle kinetics for the 6 hits are shown to the right.

Figure S4

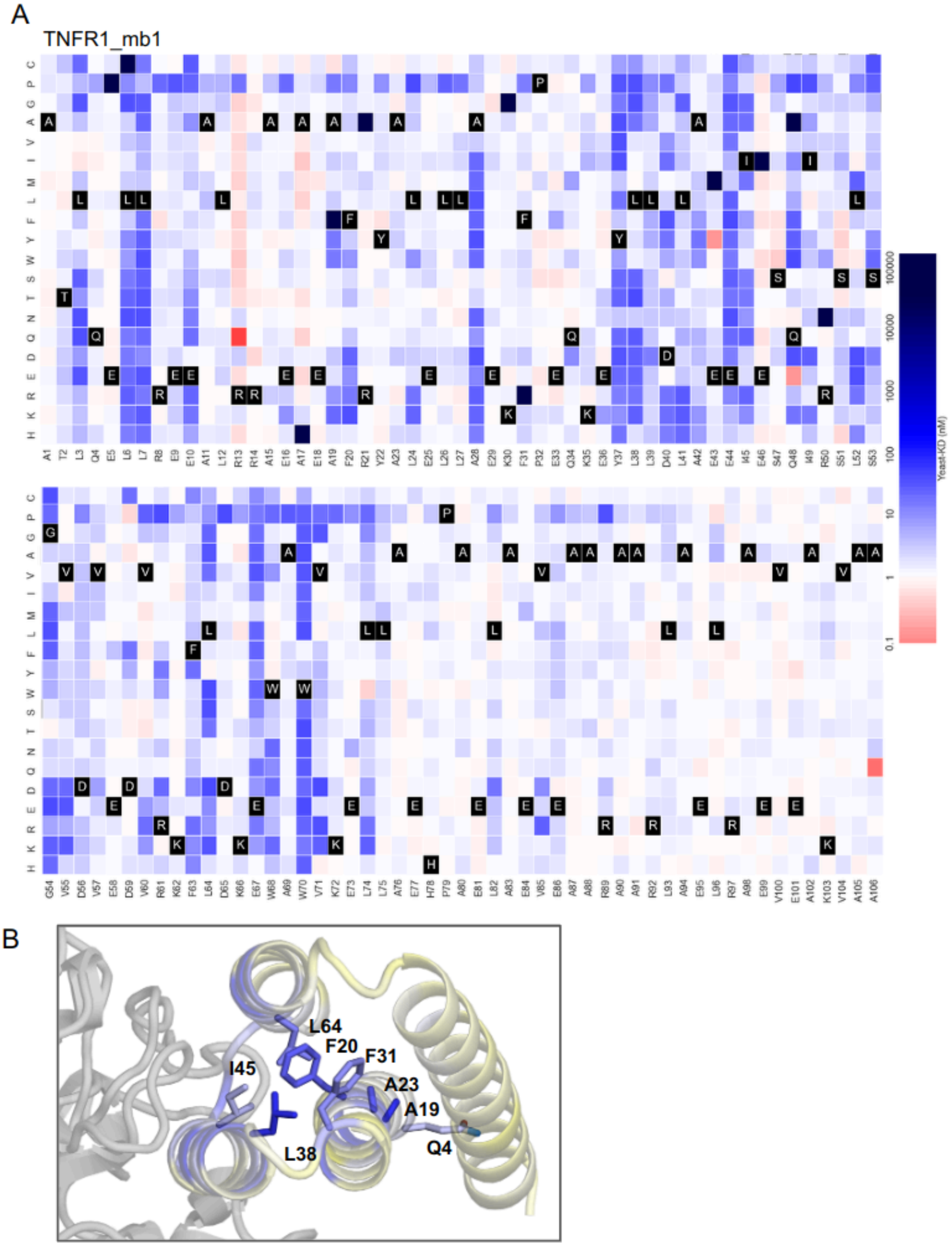


**Supplementary Fig. 4.** SSM Site saturation mutagenesis for TNFR1 binder TNFR1_mb1. A. Apparent KD in yeast display was calculated for every mutation, and is color coded, with white being similar to the original design, red indicates better binding, blue worse binding. Related to Fig. 1F. B. Positions that were strongly conserved (low entropy, blue) in the core are shown as sticks and labeled.

Figure S5

A TNFR1_mb2

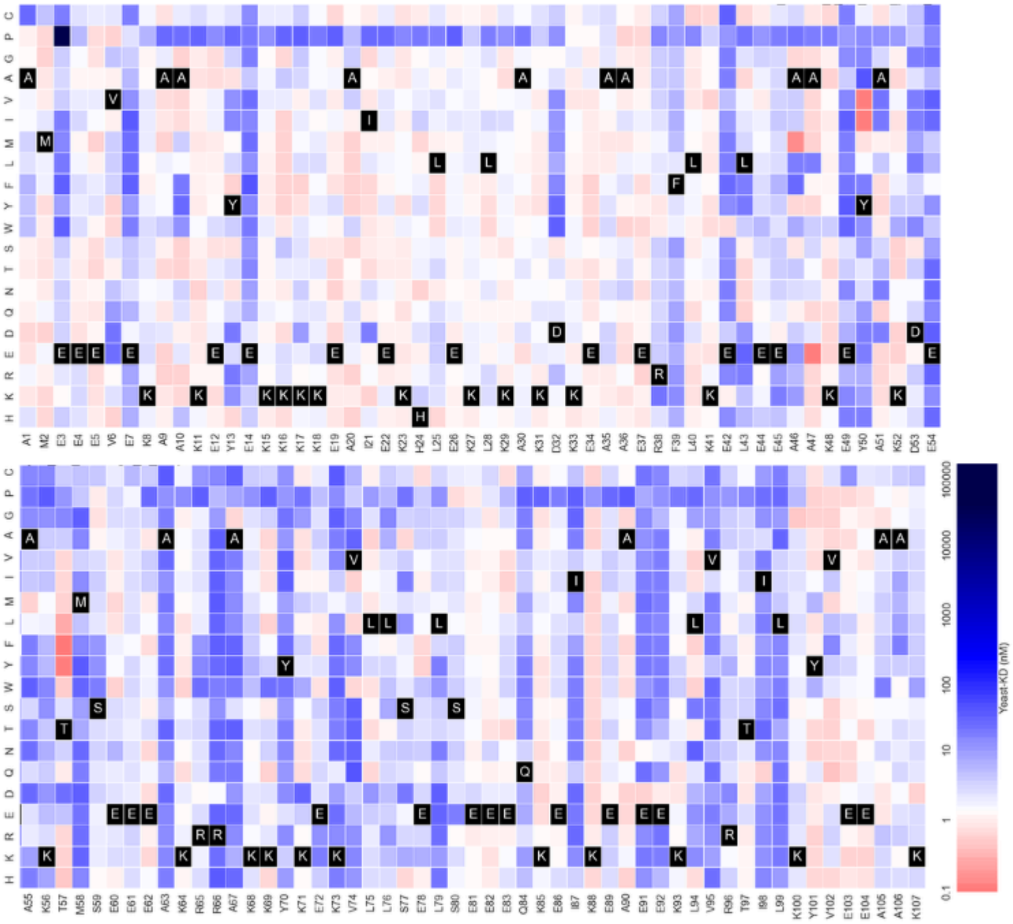


B

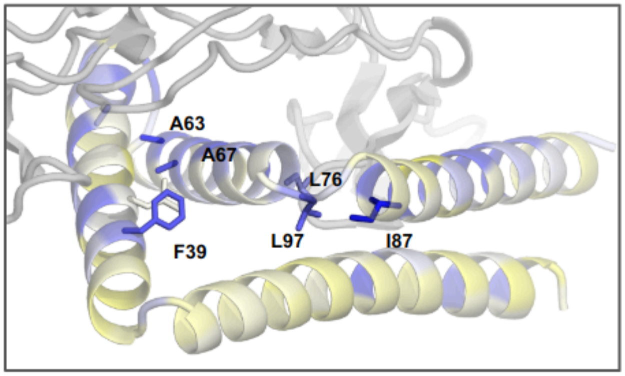


**Supplementary Fig. 5.** SSM Site saturation mutagenesis for TNFR1 binder TNFR1_mb2. A. Apparent KD in yeast display was calculated for every mutation, and is color coded, with white being similar to the original design, red indicates better binding, blue worse binding. Related to Fig. 1F. B. Positions that were strongly conserved (low entropy, blue) in the core are shown as sticks and labeled.

A

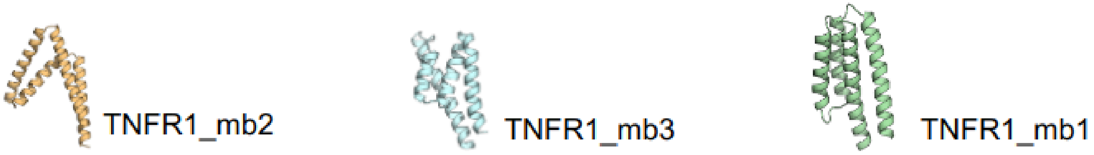


RU at 2.5uM

TNFR1_mb2

| name | TNFR 1 | mTNFR 1 | TNFR 2 | OX40 |
|---|---|---|---|---|
| MG0384 | -0.5 | 0 | -0.5 | -1 |
| MG0354 | -0.3 | -0.2 | -0.9 | -1.1 |
| MG0370 | 0 | -0.1 | -0.5 | -0.7 |
| MG0375 | 0.1 | -0.1 | -0.7 | -0.7 |
| MG0376 | 0.3 | 0.2 | -0.2 | -0.4 |
| MG0374 | 0.4 | 0.2 | -0.4 | -0.6 |
| MG0381 | 0.5 | 0.1 | -0.5 | -0.4 |
| MG0377 | 0.8 | 0.1 | -0.9 | -0.5 |
| MG0372 | 1.4 | 1.4 | 0.7 | 2.8 |
| MG0383 | 2 | 0.2 | -0.1 | -0.5 |
| MG0362 | 4.2 | 0.6 | 0 | -0.2 |
| MG0368 | 6.2 | 0.1 | -0.2 | -0.5 |
| MG0378 | 7.1 | 0.3 | 0.3 | -0.5 |
| MG0373 | 11.6 | 0.8 | 0.4 | 0.3 |
| MG0367 | 12.6 | 0.6 | -0.2 | 0 |
| MG0382 | 15.7 | 1.5 | -0.1 | -0.6 |
| MG0379 | 18.3 | 16.6 | 0 | -0.4 |
| MG0356 | 18.9 | 1.6 | 1.1 | -3.5 |
| MG0358 | 27.9 | 47.6 | -0.5 | -0.6 |
| MG0363 | 29.4 | 7.5 | 0.5 | 0.7 |
| MG0357 | 30 | 0.5 | 1.2 | -0.3 |
| MG0364 | 30.1 | 0.9 | 0.4 | 0.2 |
| MG0365 | 40.7 | 1.8 | 0.5 | 0 |
| MG0361 | 44.6 | 0.7 | -0.1 | -0.6 |
| MG0366 | 44.8 | 0.6 | 0.4 | -0.2 |
| MG0353 | 48.9 | 3.6 | 0.4 | 0 |
| MG0371 | 49 | 1 | 0.5 | 0.3 |
| MG0355 | 51.9 | 30 | 0.9 | 0.1 |
| MG0359 | 52.8 | 69 | 11.6 | -0.3 |
| MG0380 | 52.9 | 59.9 | -0.3 | -0.1 |
| MG0369 | 54.6 | 5.2 | 1 | 0.1 |
| MG0360 | 55.3 | 59.5 | 1.3 | 0.1 |

TNFR1_mb3

| name | TNFR 1 | mTNFR 1 | TNFR 2 | OX40 |
|---|---|---|---|---|
| MG0314 | -0.4 | -0.3 | -2.2 | -1.3 |
| MG0319 | -0.1 | 0 | -0.6 | -0.1 |
| MG0308 | 0.2 | 0.4 | 2.1 | -0.2 |
| MG0313 | 0.4 | 0.2 | -0.4 | -0.6 |
| MG0300 | 0.4 | 0 | -0.4 | -0.4 |
| MG0296 | 0.7 | 0.8 | -0.2 | -0.2 |
| MG0310 | 1.1 | 1.1 | 0.6 | 0.5 |
| MG0291 | 2.6 | 3.2 | 2.7 | 2.7 |
| MG0303 | 3 | 3.3 | 2.7 | 3.2 |
| MG0311 | 3 | 0.3 | -0.2 | -0.3 |
| MG0297 | 3.6 | 4.2 | 3.2 | 3.4 |
| MG0320 | 4.1 | -0.2 | -0.4 | -0.6 |
| MG0293 | 4.3 | 4.7 | 4 | 4.1 |
| MG0301 | 5.1 | 6.2 | 5.3 | 5.8 |
| MG0307 | 5.4 | 2.4 | 0.2 | 0.1 |
| MG0317 | 6.5 | 0.1 | -0.4 | 0 |
| MG0295 | 7.5 | 6.7 | 5.8 | 6.2 |
| MG0298 | 9.3 | 1 | -0.3 | -0.2 |
| MG0316 | 9.8 | -0.1 | 0 | -0.8 |
| MG0318 | 9.9 | -0.2 | -0.4 | -0.9 |
| MG0299 | 10.6 | 13.4 | 11.9 | 12.9 |
| MG0304 | 11.8 | 0.4 | -0.1 | -0.2 |
| MG0294 | 12.5 | 0.5 | -0.1 | -0.3 |
| MG0312 | 19.3 | 0.3 | -0.2 | -0.2 |
| MG0290 | 20.3 | 0.6 | 0.1 | -0.1 |
| MG0306 | 25.6 | 1.4 | -0.2 | -0.2 |
| MG0309 | 27.7 | 2.6 | 6.1 | -0.1 |
| MG0305 | 27.8 | 4.9 | 4.1 | 4.5 |
| MG0289 | 34.3 | 34.4 | 33.3 | 36.6 |
| MG0302 | 36.4 | 0.3 | 0.3 | -0.1 |
| MG0315 | 37.7 | 2.6 | -0.1 | -0.2 |
| MG0292 | 46.7 | 1 | -0.2 | -0.5 |

TNFR1_mb1

| name | TNFR 1 | mTNFR 1 | TNFR 2 | OX40 |
|---|---|---|---|---|
| MG0437 | -0.3 | -1.4 | -1.3 | -1.2 |
| MG0417 | -0.3 | -1.7 | -1.9 | -1.8 |
| MG0443 | -0.3 | -2.3 | -1.6 | -1.8 |
| MG0418 | -0.2 | -29.4 | -0.7 | -1.3 |
| MG0442 | -0.2 | -0.9 | -0.9 | -1 |
| MG0433 | -0.1 | -1.7 | -1.6 | -1.7 |
| MG0439 | -0.1 | -11.6 | -1 | -1.3 |
| MG0448 | 0 | -0.9 | -1.1 | -1 |
| MG0446 | 1 | -0.3 | 2.3 | 0.5 |
| MG0434 | 1.1 | -1.1 | -0.9 | -1.3 |
| MG0438 | 2.3 | -1 | -0.6 | -0.8 |
| MG0441 | 2.5 | -19.6 | -31.6 | -14.7 |
| MG0424 | 4.5 | -0.8 | -0.7 | -0.9 |
| MG0429 | 5 | -36.6 | -1 | -1.4 |
| MG0436 | 5.3 | -0.7 | 0.4 | -0.8 |
| MG0428 | 5.4 | -0.7 | 0.4 | -0.1 |
| MG0427 | 5.8 | -0.9 | -0.6 | -0.8 |
| MG0430 | 6.3 | -1.2 | -35 | -1.2 |
| MG0447 | 10.1 | -1.1 | -26.3 | -10.3 |
| MG0423 | 11.6 | -1.9 | -0.9 | -1.1 |
| MG0435 | 12 | -1.1 | -0.9 | -2.2 |
| MG0426 | 15.2 | -1.4 | | -0.8 |
| MG0419 | 19.2 | -1 | -0.8 | -0.9 |
| MG0444 | 24.4 | -0.5 | 1 | -0.3 |
| MG0422 | 29.1 | -1.3 | -1.1 | -1.1 |
| MG0425 | 44 | -21.7 | -0.1 | 0.1 |
| MG0432 | 49.8 | -5.7 | -0.4 | -0.4 |
| MG0420 | 52.4 | -0.4 | -0.1 | 23.9 |
| MG0431 | 54.1 | -2.4 | 30.4 | -16.9 |
| MG0421 | 54.7 | -0.9 | -0.5 | -0.7 |
| MG0445 | 56.9 | -1.8 | -0.5 | -0.7 |
| MG0440 | 80.3 | -0.8 | -0.7 | -0.9 |

B

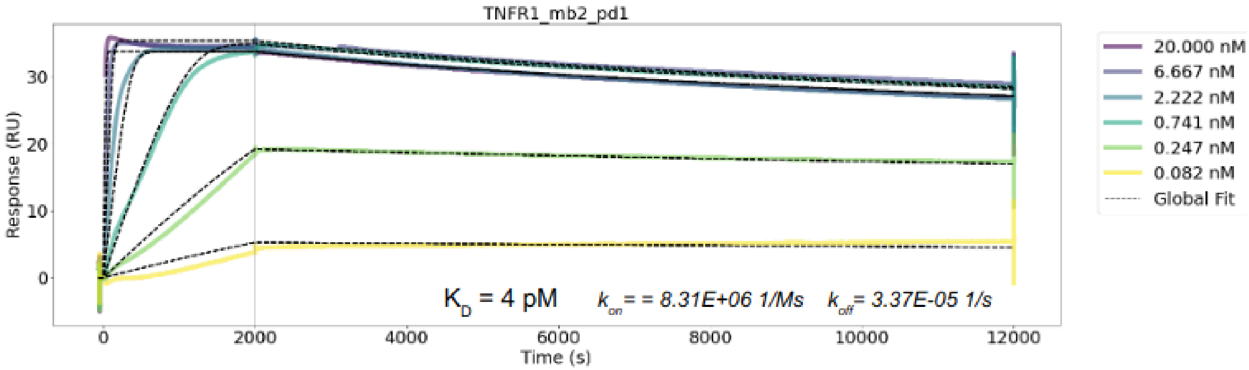


C

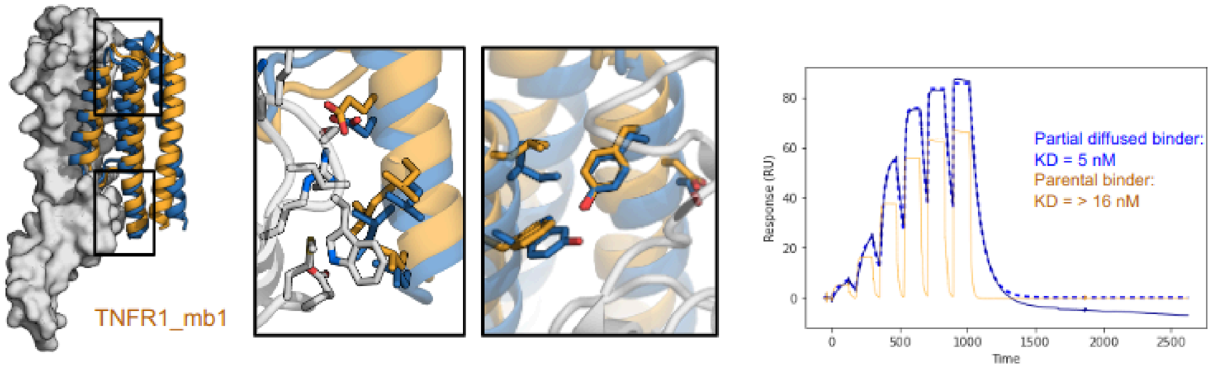


**Supplementary Fig. 6.** Partial diffusion is highly successful in generating specific binders. A. 96 partially diffused TNFR1 binders were analyzed for binding to TNFR1 and related receptors by

SPR. About 30% of variants derived from 3 starting backbones indicated above successfully retained or improved their binding. Binding was recorded as absolute response at 2.5 μM and is colorcoded, with darker purple indicating stronger binding. In most cases, no or only weak binding to related receptors TNFR2 and OX40 was recorded, while some variants gained reactivity to the mouse homolog of TNFR1 (mTNFR1). B. B. SPR traces for TNFR2_mb1, related to figure 3. $K_D$ was calculated based on global fit of $k_{on}$ and $k_{off}$. Since affinity is near instrument liments, we refer to $K_D$ as <10 pM. C. For design TNFR1_mb1 diffusion and sequence designs already found a close to ideal solution, with two of the main interfaces remaining very similar (middle inlets), and comparable affinity (SPR trace on the right) after partial diffusion.

Figure S7

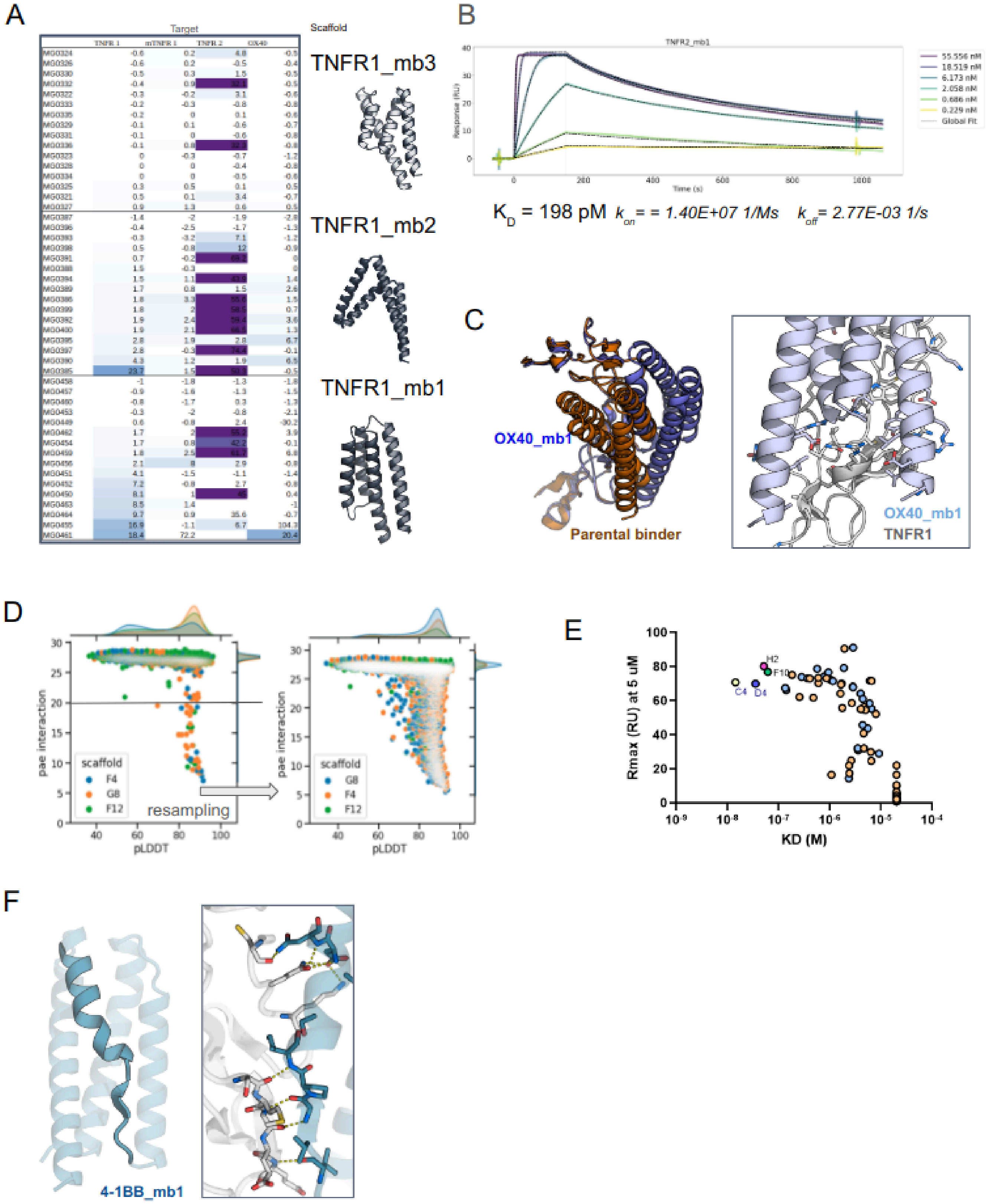

A
Target
Scaffold
TNFR1_mb3
TNFR1_mb2
TNFR1_mb1
B
TNFR2_mb1
55.556 nM
18.519 nM
6.173 nM
2.058 nM
0.686 nM
0.229 nM
Global Fit
Response (RU)
Time (s)
$K_D$ = 198 pM $k_{on}$ = = 1.40E+07 1/Ms $k_{off}$ = 2.77E-03 1/s
C
OX40_mb1
Parental binder
OX40_mb1
TNFR1
D
pae interaction
pLDDT
scaffold
F4
G8
F12
resampling
E
Rmax (RU) at 5 uM
KD (M)
H2
F10
C4
D4
F
4-1BB_mb1

**Supplementary Fig. 7.** Generation of specific binders to TNFR2, OX40 and 4-1BB

A. 48 partially diffused binders to TNFR2 were analyzed for binding to TNFR2 and related receptors by SPR. Binding was recorded as absolute response at 2.5 μM and is colorcoded, with darker purple indicating stronger binding. B. SPR traces for TNFR2_mb1, related to figure 3. $K_D$ was calculated based on global fit of $k_{on}$ and $k_{off}$ C. Comparison of backbones of the parental binder and the retargeted OX40 binder OX40_mb1. Bottom panel shows unique matching to a OX40 specific helical turn. D. Computational metric for 4-1BB directed designs after one round of partial diffusion and after a second, for which all designs that had a pae_interaction <20 in the first round were used as input. E. 48 4-1BB directed designs were analyzed by single cycle kinetics for 4-1BB binding on SPR. $K_D$ is indicated when curves could be fit, Rmax indicates the highest response on SPR at a concentration of 5 μM. F. AF2 model of a 4-1BB binder 4-1BB_mb1 which has a unique sheet and kinked helix in the interface to match the 4-1BB fold.

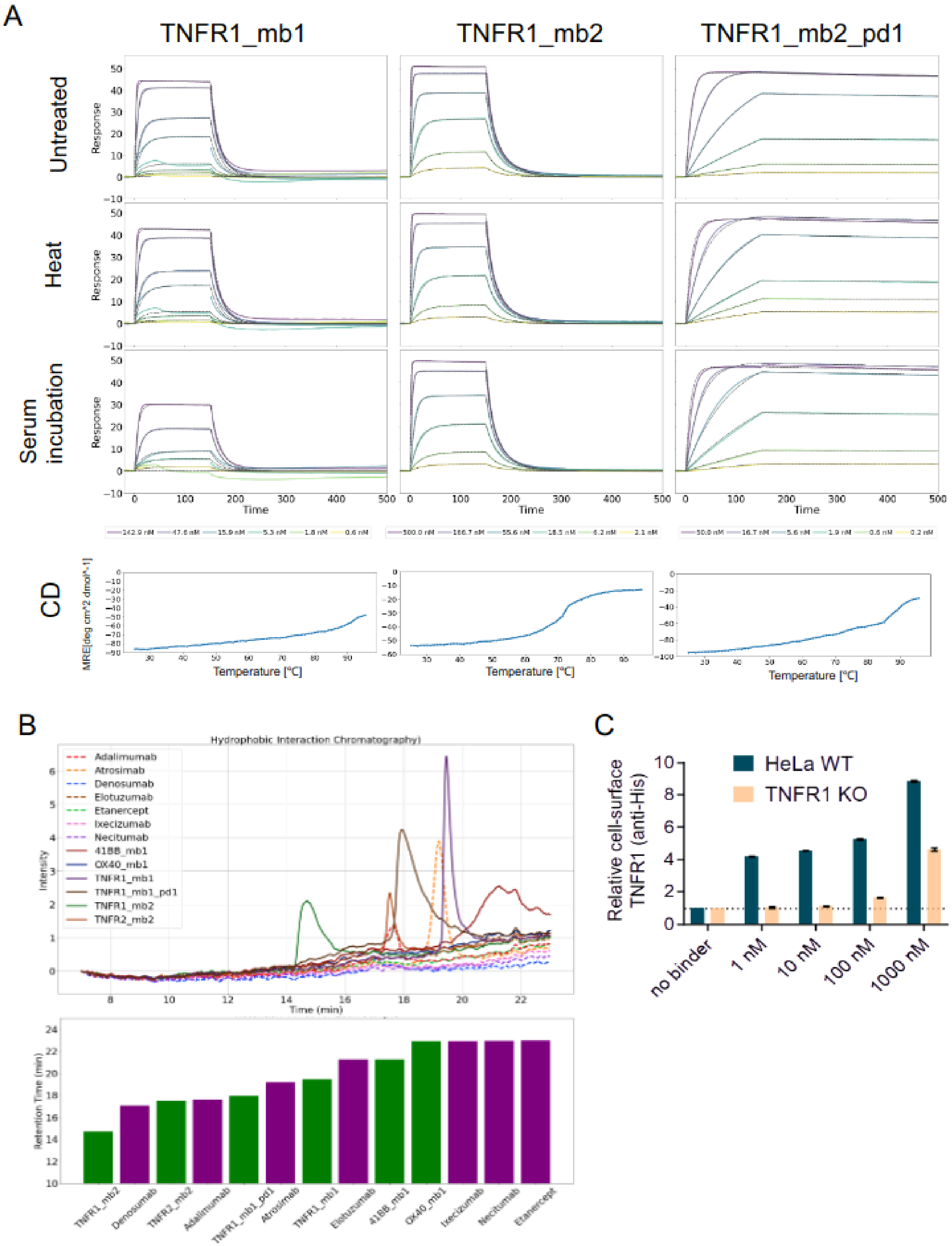


**Supplementary Fig. 8.** Stability and developability features of TNFR1 binders.

A. Stability to heat and mouse serum treatment. Binders (columns) were incubated for 15 minutes at 95°C (second row) or at a concentration of 1 mg/ml 1:1 mixed with mouse sera and incubated at 37°C for 2h (third row). SPR binding did not change substantially after this treatment. Bottom row shows unfolding as defined by change in circular dichroism signal (MRE[deg cm^2 dmol^-1]) after heating to 95°C. B. Traces and retention times during hydrophobic interaction chromatography (HIC) for minibinders and clinical molecules. Protein was loaded on a HIC column and eluted by applying a salt gradient. Longer retention indicates stronger hydrophobic interaction. C. Binding of TNFR1_mb2_pd1 to TNFR1 on HeLA cells. Cells were stained with varying concentrations of binder and signal evaluated by flow cytometry, and compared to the same cell line with TNFR1 knocked out.

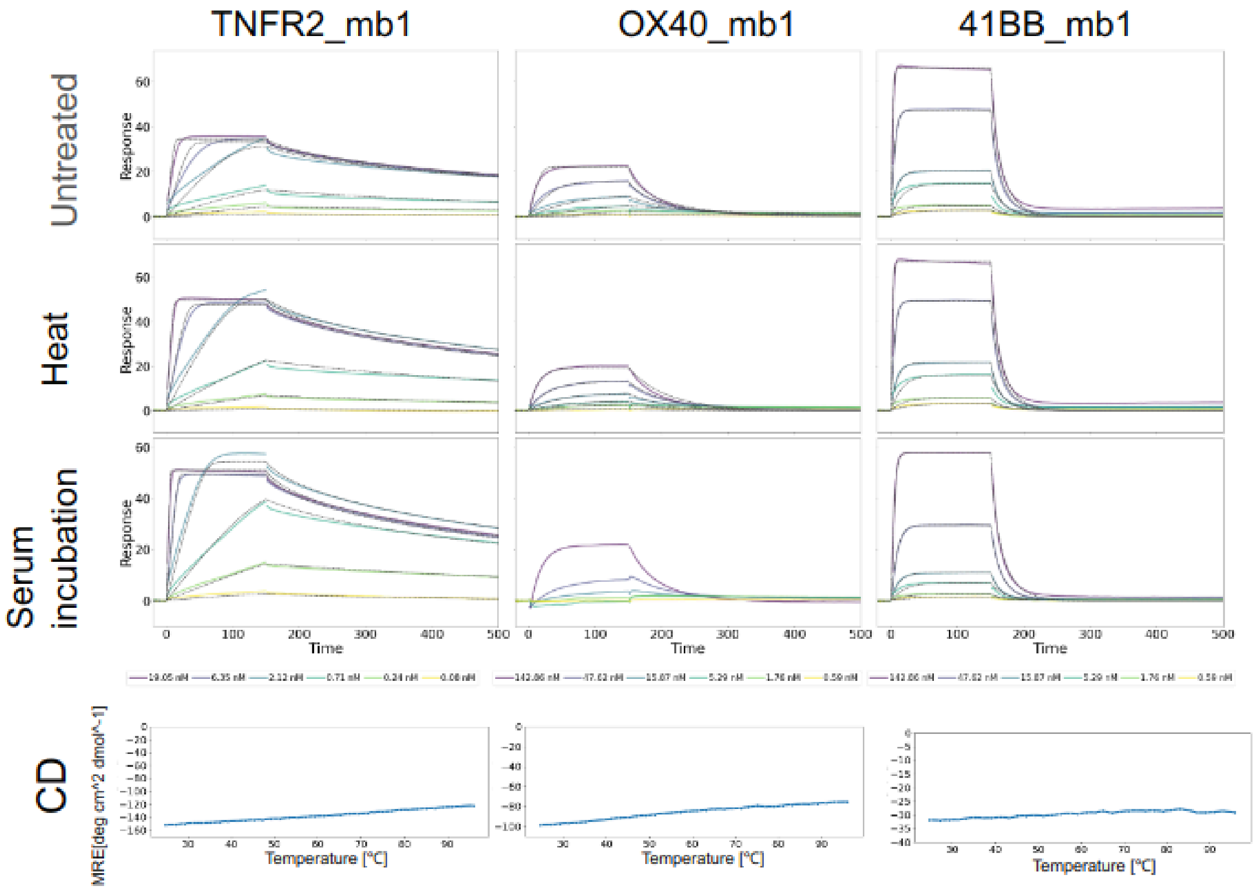


**Supplementary Fig. 9.** Stability and developability features of TNFR2, OX40 and 4-1BB binders.

Stability to heat and mouse serum treatment. Binders (columns) were incubated for 15 minutes at 95°C (second row) or at a concentration of 1 mg/ml 1:1 mixed with mouse sera and incubated at 37°C for 2h (third row). SPR binding signal did not change substantially after this treatment. Bottom row shows unfolding as defined by change in circular dichroism signal (MRE[deg cm^2 dmol^-1]) after heating to 95°C.

**Fig. S10**

**A**

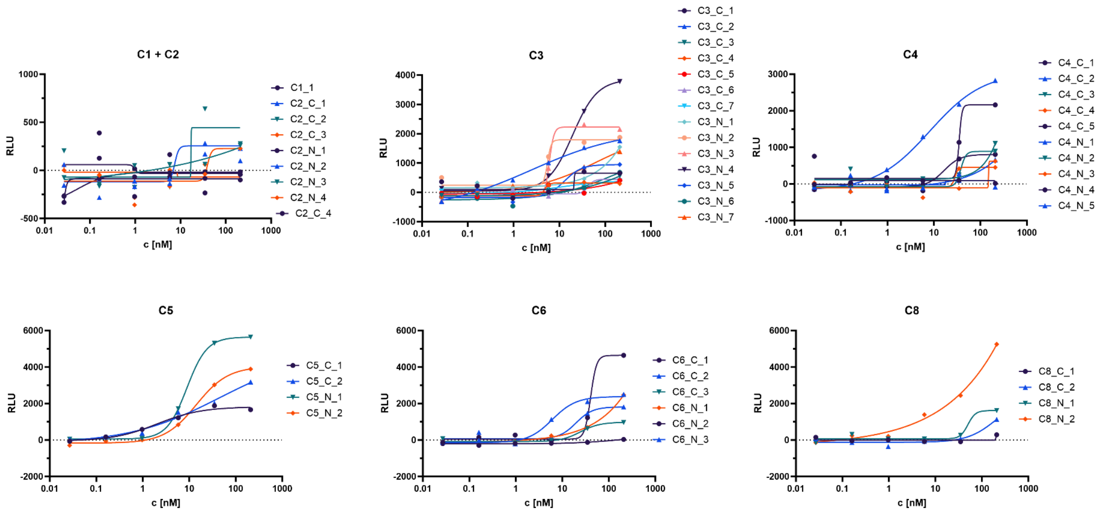


**B**

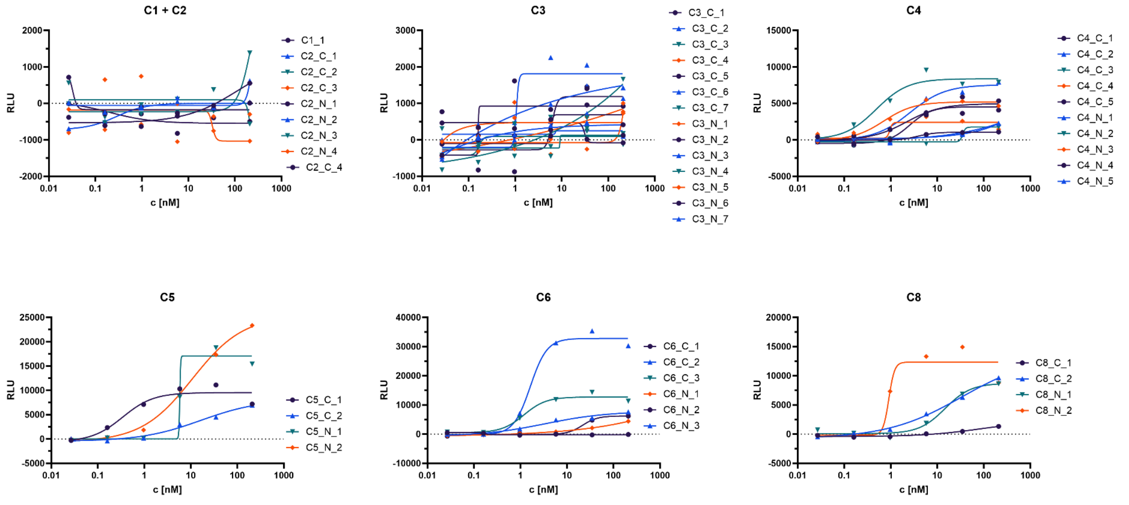


**Supplementary Fig. 10.** Agonism of 4-1BB and OX40 directed oligomerized binders.
A. Multivalent presentation of 4-1BB binder design on cyclic oligomers activates 4-1BB signaling on luciferase reporter cell lines. The curves are averaged from 2 biological replicates and show signalling activity of 46 oligomers categorized by valency. B. Multivalent presentation of OX40 binder design on cyclic oligomers activates OX40 signaling on luciferase reporter cell lines. The curves are averaged from 2 biological replicates and show signalling activity of 44 oligomers categorized by valency.

**Table 3:** Crystallographic Data Collection and Refinement Statistics

| | **TNFR2_mb1 TNFR2 complex (PDB: 9CU8)** |
|---|---|
| **Data collection** | |
| Space group | $P\ 2_1\ 2_1\ 2_1$ |
| Cell dimensions | |
| *a*, *b*, *c* (Å) | 35.07, 74.62, 104.97 |
| *α*, *β*, *γ* (°) | 90, 90, 90 |
| Resolution (Å) | 29.16 – 1.96 (2.01 - 1.96)[a] |
| $R_{merge}$ | 0.137 (0.725) |
| $I/\sigma(I)$ | 11.0 (2.60) |
| $CC_{1/2}$ | 0.997 (0.880) |
| Completeness (%) | 99.70 (95.60) |
| Redundancy | 11.0 (10.7) |
| **Refinement** | |
| Resolution (Å) | 29.16 – 1.96 |
| No. reflections | 20386 (1229) |
| $R_{\mathrm{work}}$ / $R_{\mathrm{free}}$ (%) | 0.1830/0.2199 (0.2226/0.2808) |
| No. atoms | 2363 |
| Protein | 2176 |
| Water | 187 |
| Ramachandran<br>Favored/allowed<br>Outlier (%) | 98.58/1.42<br>00.00 |
| R.m.s. deviations | |
| Bond lengths (Å) | 0.006 |
| Bond angles (°) | 0.799 |
| $B_{\mathrm{factors}}$ ($Å^2$) | |
| Protein | 30 |
| Water | 35 |

1. Data were collected from one crystal per condition.
2. [a]Values in parentheses are for the highest-resolution shell.
